# Pathological Tau transmission initiated by binding lymphocyte-activation gene 3

**DOI:** 10.1101/2023.05.16.541015

**Authors:** Chan Chen, Ramhari Kumbhar, Hu Wang, Xiuli Yang, Kundlik Gadhave, Cyrus Rastegar, Yasuyoshi Kimura, Adam Behensky, Sruthi Katakam, Deok Jeong, Liang Wang, Anthony Wang, Rong Chen, Shu Zhang, Lingtao Jin, Creg J. Workman, Dario A.A. Vignali, Olga Pletinkova, David W. Nauen, Philip C. Wong, Juan C. Troncoso, Mingyao Ying, Valina L. Dawson, Ted M. Dawson, Xiaobo Mao

## Abstract

The spread of prion-like protein aggregates is believed to be a common driver of pathogenesis in many neurodegenerative diseases. Accumulated tangles of filamentous Tau protein are considered pathogenic lesions of Alzheimer’s disease (AD) and related Tauopathies, including progressive supranuclear palsy, and corticobasal degeneration. Tau pathologies in these illnesses exhibits a clear progressive and hierarchical spreading pattern that correlates with disease severity^1, 2^. Clinical observation combined with complementary experimental studies^3, 4^ have shown that Tau preformed fibrils (PFF) are prion-like seeds that propagate pathology by entering cells and templating misfolding and aggregation of endogenous Tau. While several receptors of Tau are known, they are not specific to the fibrillar form of Tau. Moreover, the underlying cellular mechanisms of Tau PFF spreading remains poorly understood. Here, we show that the lymphocyte-activation gene 3 (Lag3) is a cell surface receptor that binds to PFF, but not monomer, of Tau. Deletion of *Lag3* or inhibition of Lag3 in primary cortical neurons significantly reduces the internalization of Tau PFF and subsequent Tau propagation and neuron-to-neuron transmission. Propagation of Tau pathology and behavioral deficits induced by injection of Tau PFF in the hippocampus and overlying cortex are attenuated in mice lacking *Lag3* selectively in neurons. Our results identify neuronal Lag3 as a receptor of pathologic Tau in the brain, and for AD and related Tauopathies a therapeutic target.

**One Sentence Summary:** Lag3 is a neuronal receptor specific for Tau PFF, and is required for uptake, propagation and transmission of Tau pathology.

## Main

In Alzheimer’s disease (AD), the burden of Tau pathology is highly correlated with the severity of cognitive decline^5, 6^. Emerging postmortem and experimental studies indicate that pathogenic Tau spreads in a stereotypical manner, driving disease progression^1–4, 7, 8^. Pathogenic Tau preformed fibrils (PFF) serve as prion seeds that induce the misfolding of endogenous tau monomer, resulting in cell-to-cell propagation in Tauopathies^3, 4, 7^. Recent work has identified several receptors of Tau uptake, including heparan sulfate proteoglycans (HSPGs)^9–12^ and low-density lipoprotein receptor-related protein 1 (LRP1)^13^. However, these identified receptors are not specific to Tau aggregates or other Tau species (e.g. monomer and soluble Tau)^9–13^. Of note, substantial amount of Tau PFF still accumulates in LRP1 knocked down cells^13^, which suggests that there are other receptors or pathways that could mediate the endocytosis of Tau PFF. Identification of a Tau-fibril-specific receptor could provide the opportunity to validate its role in initiating Tauopathy and offer a potential therapeutic target for AD and related Tauopathies.

While several Tau receptors (e.g. HSPGs, LRP1) have shown to bind α-synuclein (α-syn)^9, 14^, we identified lymphocyte-activation gene 3 (Lag3) as a major cell-surface receptor that specifically binda with α-syn fibrils, but not to the α-syn monomer^15, 16^. Depletion of *Lag3* inhibits the uptake and reduces the subsequent α-syn pathology^15–18^. To determine whether Lag3 is specific for α-syn fibrils, but not to other prion-like proteins, we further tested the interaction between Lag3 and tau PFF. The results showed no interaction between Lag3 and heparin-induced Tau (hep-Tau) PFF^15, 16^. A recent study showed that hep-Tau PFF is not an active seed in vivo, as compared to self-seeded tau or the AD brain extracts-induced Tau (AD-Tau) PFF^3^. Thus, we wondered whether bioactive tau, but not hep-tau, fibrils binds to Lag3. Here we show that Lag3 is a cell surface receptor that is specific for pathogenic fibrils, but not the monomer, of Tau. We employed both genetic and biochemical approaches to test the requirement of Lag3 for pathogenic Tau propagation and neuron-to-neuron transmission.

### Preparation and characterization of Tau PFF

Intracerebral inoculation of Tau PFF purified from AD brains (AD-Tau) resulted in robust Tau inclusions in anatomically connected brain regions in non-transgenic mice^3, 4^. We extracted AD-Tau, seeded recombinant Tau (5/95%) for fibrillization, and sonicated to obtain Tau PFF as published^3^. We characterized the Tau-PFF with transmission electron microscopy (TEM) (Extended Data Fig. 1a-b) and thioflavin T (ThT) assay (Extended Data Fig. 1c-d).

We labeled Tau with biotin to distinguish it from endogenous Tau, and then investigated the interaction between Tau-biotin species and primary cortical neurons using a cell surface binding assay. Tau-biotin PFF binds to primary cortical neurons as detected with streptavidin-AP (alkaline phosphatase) staining^19–21^ (Extended Data Fig. 2a-b). Tau-biotin PFF binding to neurons is saturable with an apparent dissociation constant (KD) of 276 nM, whereas both hep-Tau-biotin PFF and Tau-biotin monomer fail to show the saturable binding (KD > 3000 nM) (Extended Data Fig. 2a-b). Biotin as the negative control showed a minimal binding signal (Extended Data Fig. 2a-b) suggesting the existence of PFF-specific receptor(s) and/or binding site(s) for Tau PFF. These results further suggest that heparin can reduce the interaction between neuronal surface and Tau-biotin PFF.

### The Interaction between Lag3 and Tau-biotin PFF

In the well-established cell surface binding assay^19–21^, Tau-biotin PFF at the indicated concentrations were administered individually into the Lag3-expressing SH-SY5Y cells, and the binding signals as detected with streptavidin-AP and its substrate staining. Our results showed that Tau-biotin PFF binds to Lag3 in a saturable manner, with a KD = 158 nM, whereas no signal was observed with the Tau-biotin monomer (Fig. 1a-b, Extended Data Fig. 3a-b).

**Figure 1:**
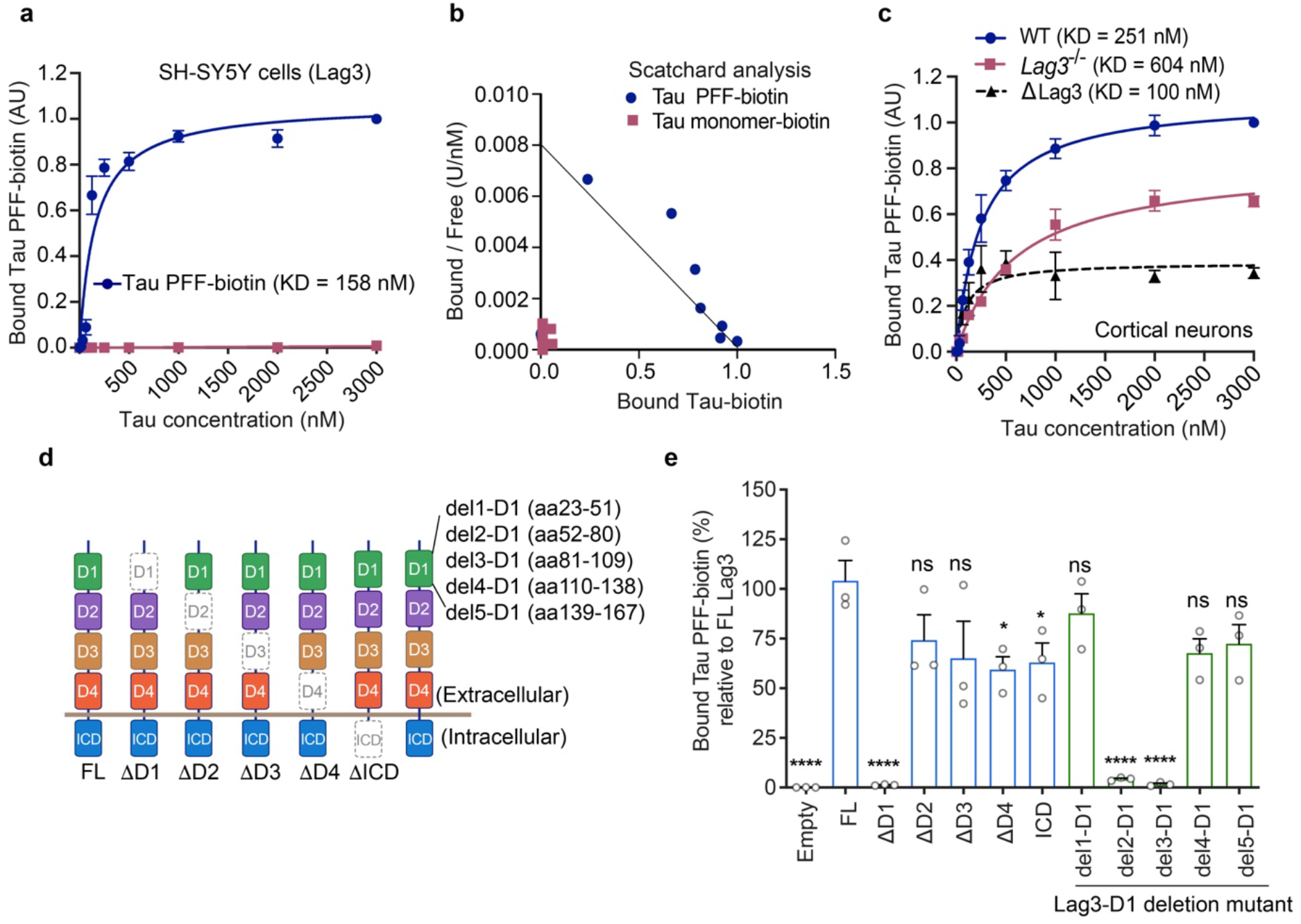
Lag3 binds with Tau PFF, but not Tau monomer. **a.** Tau-biotin PFF binds to Lag3-expressing SH-SY5Y cell surface in a saturable manner, as a function of Tau-biotin total concentration (monomer equivalent for PFF preparations). **b.** Scatchard analysis. Data are the means ± SEM, *n* = 3 independent experiments. **c.** Binding of Tau-biotin PFF to mouse primary cortical neurons is reduced by *Lag3* knockout (*Lag3*^-/-^). Tau-biotin PFF WT-KD = 251 nM, *Lag3*^−/−^-KD = 604 nM, estimated KD for neuronal Lag3 (dotted line, ΔLag3 = wild type minus *Lag3*^−/−^) is 100 nM. Data are the means ± SEM, *n* = 3 independent experiments. **d.** Schematics of deletion mutants of Lag3 ectodomains. **e.** Tau-biotin PFF binding to Lag3 and deletion mutants of Lag3: extracellular domains (ΔD1-ΔD4), intracellular domain (ΔICD, and subdomains of D1 domain (del1—5-D1) (*n* = 3). \*\*\*\**P* < 0.05, \*\*\*\**P* < 0.0001 compared to FL, n.s., not significant, one-way ANOVA followed by Dunnett’s correction. Data are the means ± SEM.

We have determined that Lag3 is expressed in neurons using two in vivo systems^18^. Thus, we further studied the interaction between Lag3 and pathogenic Tau in neurons. WT mouse cortical neurons demonstrated Tau-biotin PFF-binding (KD = 251 nM), whereas *Lag3*^-/-^ mouse cortical neurons had reduced Tau-biotin PFF-binding (Fig. 1c, Extended Data Fig. 3c-d). Specific binding of Tau-biotin PFF to Lag3 in primary cortical neurons was determined by subtracting the binding in WT neuron cultures from the binding in *Lag3*^-/-^ neuron cultures, and reveals a KD of 100 nM (Fig. 1c). These results taken together show that Tau-biotin PFF specifically binds to Lag3, but not to Tau-biotin monomer.

Heparin can inhibit cellular uptake of Tau^9^ by mimicking HSPGs binding with Tau. We further performed the cell surface binding assay to compare hep-Tau-biotin PFF and AD-Tau-induced Tau-biotin PFF. The result is consistent with the previous work^15^ that hep-Tau-biotin PFF exhibited significantly less binding to *Lag3*-transfected SH-SY5Y cells (Extended Data Fig. 4a-b), compared to AD-Tau-induced Tau PFF-biotin.

Lag3 has an ectodomain composed of four immunoglobulin (Ig)-like domains (D1-D4)^22^. To determine the Tau-biotin PFF-binding domain, we sequentially deleted each Ig-like domain of Lag3 and performed the cell surface binding assay with overexpression of Lag3 deletion mutants. These experiments revealed that Tau-biotin PFF preferentially binds to the D1 domain, and deletion of the D2, D3, D4 or intracellular domain (ICD) reduced the binding to some extent (Fig. 1d-e, Extended Data Fig. 5a-b). As deletion of the D1 can eliminate the Tau-biotin PFF-binding to Lag3, additional deletions of D1 subdomains (del1—5) were examined. Del2-D1 and Del3-D1 can abolish the Tau-biotin PFF-binding to Lag3, whereas del1-D1, del4-D1, and del5-D1 did not show significantly reduced binding to Lag3 (Fig. 1e, Extended Data Fig. 5a,c). These results show that residues 52-109 in the D1 domain are essential for the Lag3 interaction with Tau-biotin PFF.

### Neurons uptake Tau PFF via Lag3

To determine whether Lag3 is involved in the endocytosis of Tau PFF, pHrodo red dye was conjugated with Tau PFF, so that Tau-pHrodo PFF can fluoresce as pH decreases from the neutral cytosolic pH to the acidic pH of the endosome and lysosome^23^. Tau-pHrodo PFF fluoresced and the fluorescent signal was increased in WT cortical neurons with time via live imaging, whereas *Lag3*^-/-^ neurons exhibited minimal fluorescent signal (Fig. 2a-b). Overexpression of Lag3 by lentiviral transduction in WT neurons enhanced the fluorescent signal, and overexpression of Lag3 in *Lag3*^-/-^ neurons restored the uptake of Tau-pHrodo PFF (Fig. 2a-b). These results show that neuronal Lag3 can uptake Tau-pHrodo PFF.

**Figure 2:**
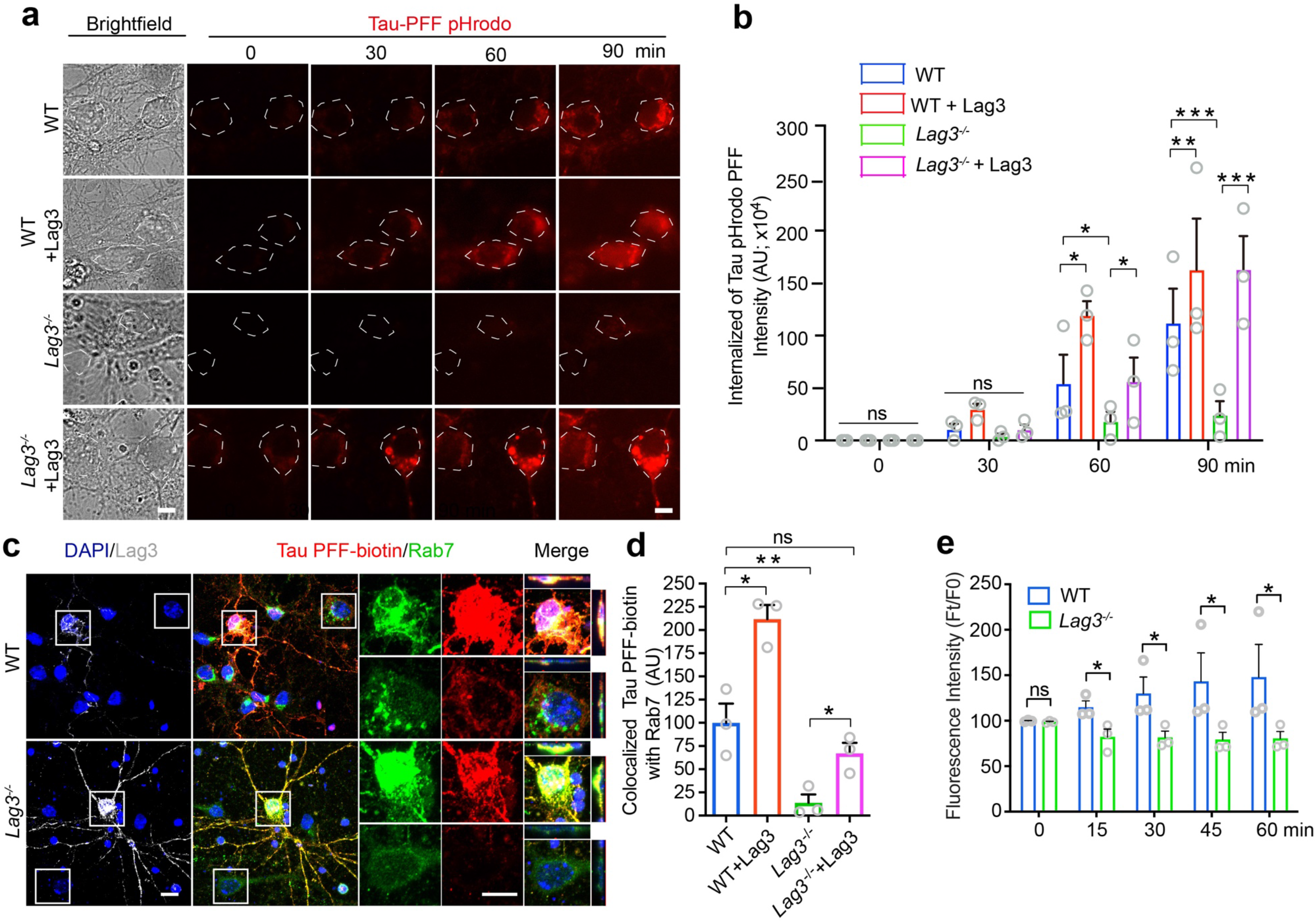
Lag3 is required for neuronal uptake of Tau PFF. **a.** Live image analysis of the endocytosis of Tau PFF-pHrodo in WT, WT+Lag3, *Lag3*^-/-^, and *Lag3*^-/-^+Lag3 neurons at indicated time points. Increased red fluorescence indicates higher endocytosis. Scale bar, 10 µm **b.** Quantification of Tau PFF-pHrodo, *n* = 3 independent experiments. **c.** *Lag3*^-/-^ reduced the neuronal uptake of Tau-biotin PFF co-localized with endosome marker (Rab7). WT or *Lag3*^-/-^ neurons were cultured, and received transient transfection with Lag3. WT, WT+Lag3, *Lag3*^-/-^, and *Lag3*^-/-^+Lag3 neurons were treated with 500 nM Tau-biotin PFF for 2 hr. Scale bar, 10 µm. **d.** Quantification of c from 3 independent experiments. One-way analysis of variance (ANOVA) with Tukey’s correction. **e.** Quantification of time series imaging of calcium-dependent fluorescence of WT and *Lag3*^-/-^ neurons. Neurons were treated with 1 µM Fluo-2 acetoxymethyl (AM) ester for 30 min followed by 500 nM Tau PFF and live images were acquired for the indicated time period. Error bars represent SEM. \**P* < 0.05, \*\**P* < 0.01, \*\*\**P* < 0.001.

Rab7 is a small guanosine triphosphatase (GTPase) that belongs to the Rab family and controls transport to the late endosome and lysosomes^24, 25^. We used Rab7 as an endocytosis marker to assess the co-localized signal with Tau-biotin PFF as an additional monitor of uptake. We found that Tau-biotin PFF was co-localized with Rab7 in WT cortical neurons, and that there was a significantly reduced co-localized signal in *Lag3*^-/-^ cortical neurons (Fig. 2c,d). Overexpression of Lag3 in WT and *Lag3*^-/-^ neurons enhanced the intensity of Tau-biotin PFF co-localizing with Rab7 (Fig. 2c,d).

Because calcium is involved in Tau pathogenesis^26^, calcium influx was monitored in response to Tau PFF. Perfusion of Tau PFF (500 nM) onto WT cortical neurons caused a gradual increase in intracellular calcium level (Fig. 2e, Extended Data Fig. 6). In contrast, a significant reduction of Tau PFF-induced calcium influx was observed in *Lag3*^-/-^ cortical neurons (Extended Data Fig. 6).

Taken together, these data indicate that after Tau PFF administration, Lag3 is required for the neuronal uptake of Tau PFF, and deletion of *Lag3* can reduce the subsequent calcium influx induced by Tau PFF.

### Deletion of *Lag3* attenuates Tau PFF-induced pathology

To determine the role of Lag3 in mediating Tau pathology, we administered Tau PFF in WT and Lag3^-/-^ primary cortical neurons on 7 DIV (days in vitro) and incubated for a further 14 days, and then assessed the Tau pathology, including the insoluble Tau and hyper-phosphorylated Tau (p-Tau). We fixed the neurons with methanol to remove soluble Tau as previously published^3^, and immunostained with T49, a mouse Tau-specific monoclonal antibody. The results show a significant increase of insoluble Tau in WT neurons treated with Tau PFF, compared to PBS-treated neurons (Fig. 3a-b). Depletion of *Lag3* significantly reduced the insoluble Tau (∼80%) compared to WT neurons (Fig. 3a-b). Overexpression of Lag3 by lentiviral transduction resulted in a significant increase of insoluble Tau in WT and *Lag3*^-/-^ neurons (Fig. 3a-b). Furthermore, we assessed the p-Tau level with AT8 immunostaining using anti-p-Tau (Ser202, Thr205). The results show a substantial p-Tau signal in WT neurons induced by Tau PFF as published^3^. In contrast, 75% of p-Tau signal was decreased in *Lag3*^-/-^ neurons (Fig. 3c-d). Overexpression of Lag3 significantly enhanced p-Tau signal in WT neurons and restored p-Tau signal in *Lag3*^-/-^ neurons (Fig. 3c-d).

**Figure 3:**
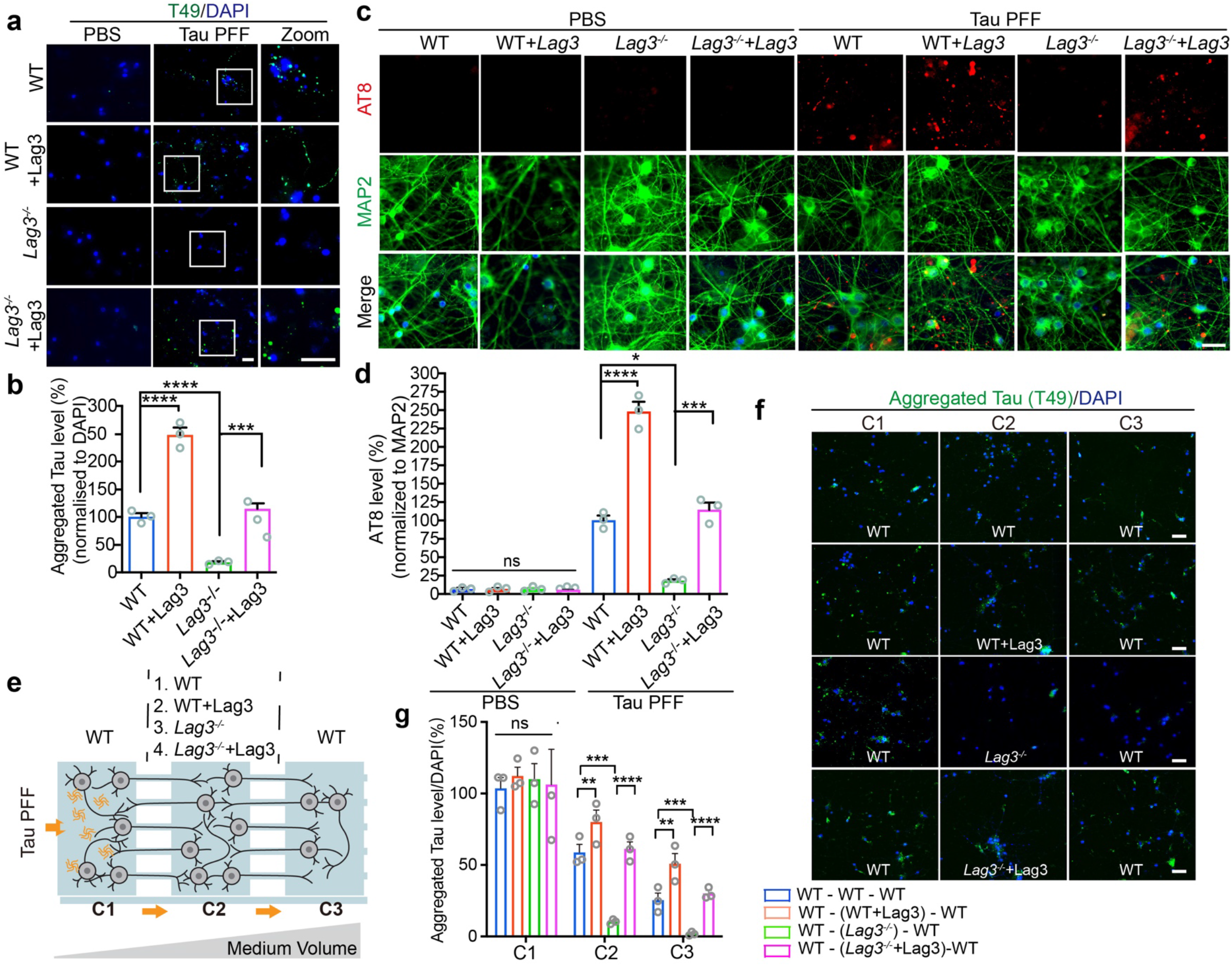
Lag3 is required for Tau pathology propagation and neuron-to-neuron transmission. **a.** Depletion of Lag3 reduces abnormal Tau aggregation (T49). WT, WT+Lag3, *Lag3*^-/-^, and *Lag3*^-/-^+Lag3 primary cortical neurons at 7 DIV were treated with Tau PFF or PBS for 14 days. Neurons were fixed with methanol to remove soluble Tau as described^3^, and immunostained with T49, a mouse Tau-specific monoclonal antibody. Abnormal Tau aggregates were assessed by immunostaining with T49 antibody. Scale bar, 20 µm. **b.** Quantification of a, *n* = 3 independent experiments. Error bars represent SEM. Statistical significance was determined using one-way ANOVA followed with Tukey’s correction, **c.** Depletion of *Lag3* reduced hyper-phosphorylated Tau at Ser202, Thr205 (AT8). Scale bar, 20 µm. **d.** Quantification of panel c, n = 3 independent experiments. Error bars represent SEM. Statistical significance was determined using one-way ANOVA followed with Tukey’s correction. **e.** Schematic of microfluidic neuron device with three chambers (chamber 1-2-3, C1-C2-C3), to separate neuron cultures. **f.** Transmission of aggregated Tau, 14 days after the administration of Tau PFF in C1. Different neurons in C2, listed as C1-(C2)-C3, are: WT-(WT)-WT, WT-(WT+Lag3)-WT, WT-(*Lag3*^−/−^)-WT, WT-(*Lag3*^−/−^+Lag3)-WT. Scale bar, 100 μm. **g.** Quantification of f. error bar represents SEM, *n* = 3 independent experiments. Statistical significance was determined using one-way ANOVA followed by Tukey’s correction, \**P* < 0.01, \*\**P* < 0.01, \*\*\**P* < 0.001, \*\*\*\**P* < 0.0001, n.s., not significant.

### Neuron-to-neuron transmission of Tau pathology is reduced in *Lag3*^-/-^ neurons

To examine the neuron-to-neuron transmission of Tau pathology, we used as microfluidic device (three separated chambers connected in tandem by two series of microgrooves) as previously described^15, 18, 27^ (Fig. 3e). Four experimental and control groups with primary cortical neurons were cultured in successive chambers (C1)-(C2)-(C3): (WT)-(WT)-(WT), (WT)-(WT+Lag3)-(WT), (WT)-(*Lag3*^-/-^)-(WT), and (WT)-(*Lag3*^-/-^+Lag3)-(WT). Using this system, the neuron-to-neuron transmission of Tau pathology was monitored via anti-T49 (insoluble Tau). In (WT)-(WT)-(WT) cortical neuron cultures, Tau PFF was administered into C1, which led to Tau pathology transmission to C2 and later C3 to (Fig. 3f). In the C1 of (WT)-(Lag3^-/-^)-(WT) cortical neuron cultures, there is no significant difference of Tau pathology compared to C1 of (WT)-(WT)-(WT), which indicates that the levels of pathogenic Tau from WT neurons (donor cells) were similar. Of note, a significant reduction of insoluble Tau can be observed in *Lag3*^-/-^ neurons in C2, which further prevented Tau propagation in WT neurons in C3 (Fig. 3f-g). Overexpression of Lag3 by lentiviral-transduction in C2 of (WT)-(WT+Lag3)-(WT) and (WT)-(*Lag3*^-/-^+Lag3)-(WT) can significantly increase the neuron-to-neuron transmission of Tau pathology in C2 and C3 (Fig. 3f-g). These results show that Lag3 is required for the neuron-to-neuron transmission of Tau pathology.

### Antibodies to Lag3 block neuronal Tau propagation and neuron-to-neuron transmission induced by Tau PFF

To determine whether Lag3 can serve as a therapeutic target, we administered Lag3 antibodies into primary cortical neurons. Lag3 antibody 410C9^28^ (30 nM) significantly blocked the binding of Tau-biotin PFF (250 nM) to Lag3-overexpressing SH-SY5Y cells, compared to the control IgG (Fig. 4a-b). Similarly, results were likewise shown by other Lag3 antibodies C9B7W^29^ (30 nM) (Extended Data Fig. 7a-b) and 17B4 (30 nM) (Extended Data Fig. 7c-d) for human LAG3. Furthermore, 410C9 reduced the Tau PFF-induced p-Tau in WT cortical neurons, compared to the control IgG (Fig. 4c-d), and C9B7W showed a similar efficacy (Extended Data Fig. 7e-f). In the in vitro assay of neuron-to-neuron transmission of Tau pathology, treatment of 410C9 antibody in C2 can significantly block Tau pathology transmission in C2 and C3 of WT cortical neurons, compared to control IgG treatment (Fig. 4e-g). Thus, these Lag3 antibodies studies support the notion that Lag3 is required for propagation and transmission of Tau pathology in vitro.

**Figure 4:**
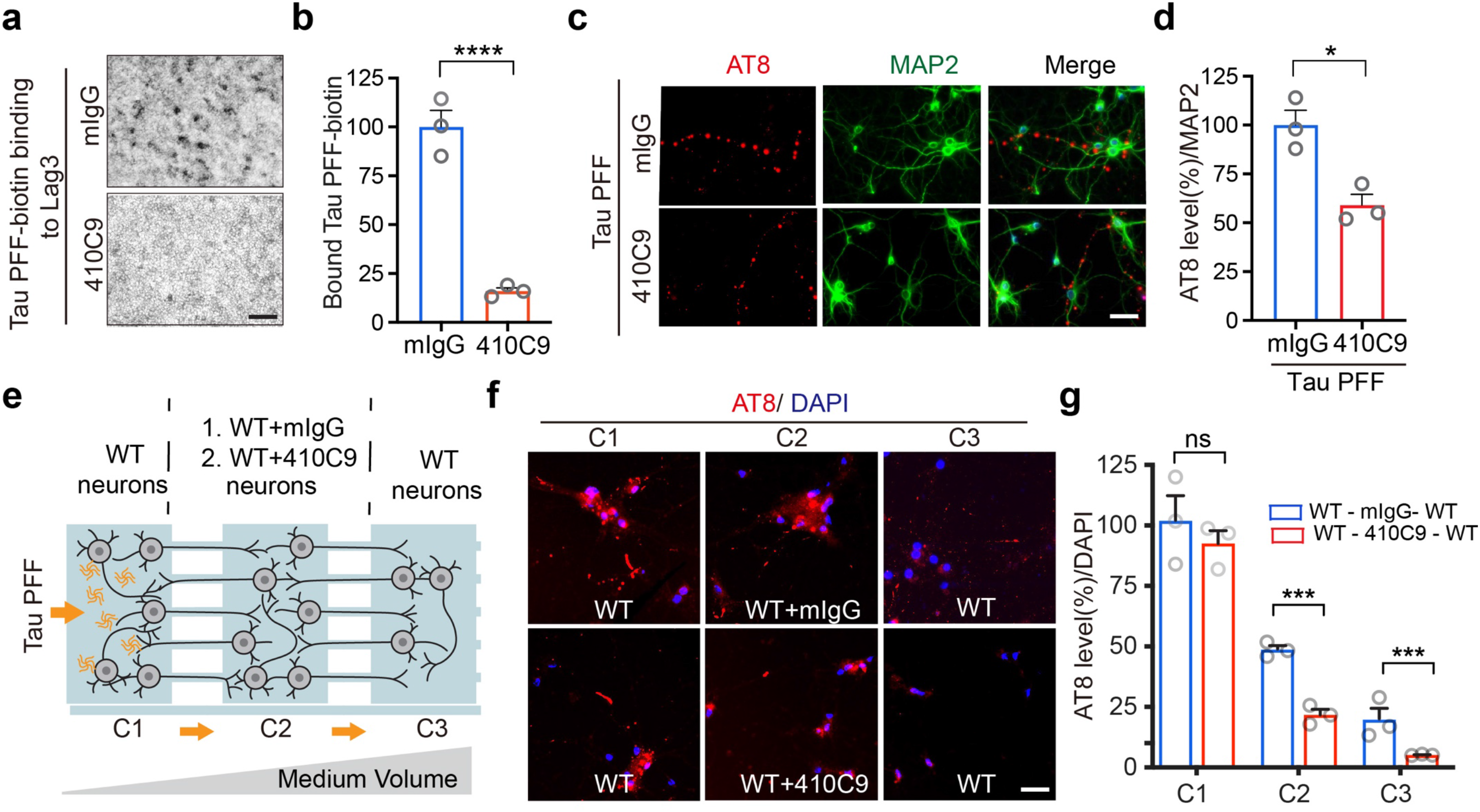
Lag3 antibodies block Tau PFF binding to Lag3, and subsequent pathologic propagation and neuron-to-neuron transmission. **a.** Anti-Lag3 410C9 blocks the binding of Tau-biotin PFF to Lag3-expressing SH-SY5Y cells. **b.** Quantification of a. Error bars represent means ± SEM, *n* = 3 independent experiments, Student’s *t*-test. \*\*\*\**P* < 0.0001. **c.** AT8 phosphorylated Tau (P-Tau) was reduced by 410C9 (30 nM) in primary cortical neurons. **d.** Quantification of c. Error bars represent means ± SEM, *n* = 3 independent experiments, Student’s *t*-test. \**P* < 0.05, Scale bar, 50 μm **e.** Schematic of a microfluidic device experimental design. The different combinations of neurons in C2, listed as C1-(C2)-C3, are: WT-(WT+mIgG)-WT, WT-(WT+410C9)-WT. **f.** Transmission of Tau pathology 14 days after the administration of Tau PFF in C1. 410C9 significantly reduces the neuron-to-neuron transmission of Tau pathology (P-Tau). **g.** Quantification of f. Error bars represent means ± SEM, *n* = 3 independent experiments, Student’s *t*-test. \*\*\**P* < 0.001, n.s., not significant. Scale bar, 50 μm.

### Deletion of *Lag3* reduces Tau pathology transmission and alleviates behavioral deficits in vivo

To determine whether neuronal Lag3 is necessary for Tau pathology transmission in vivo, we bred *Lag3*^L/L-YFP^ conditional knockout-reporter mice^30^ (*Lag3*^L/L-YFP^) with nestin^Cre^ mice (Jax Lab, strain: 003771)^31, 32^ to generate the Lag3 neuronal conditional knockout (*Lag3*^L/L-N-/-^) mice. We stereotactically injected Tau PFF into the dorsal hippocampus of *Lag3*^L/L-N-/-^ and *Lag3*^L/L-YFP^, with PBS serving as the control^3^ (Fig. 5a). Nine months after the injection of Tau PFF, p-Tau immunoreactivity was monitored in the ventral hippocampus (VH), entorhinal cortex (EC), and amygdala (Amy) (Fig. 5b-g). Substantial p-Tau signal can be found in *Lag3*^L/L-YFP^ mice, while p-Tau signal was significantly reduced in the VH (Fig. 5b-c), EC (Fig. 5d-e), and Amy (Fig. 5f-g) of *Lag3*^L/L-N-/-^ mice.

**Figure 5:**
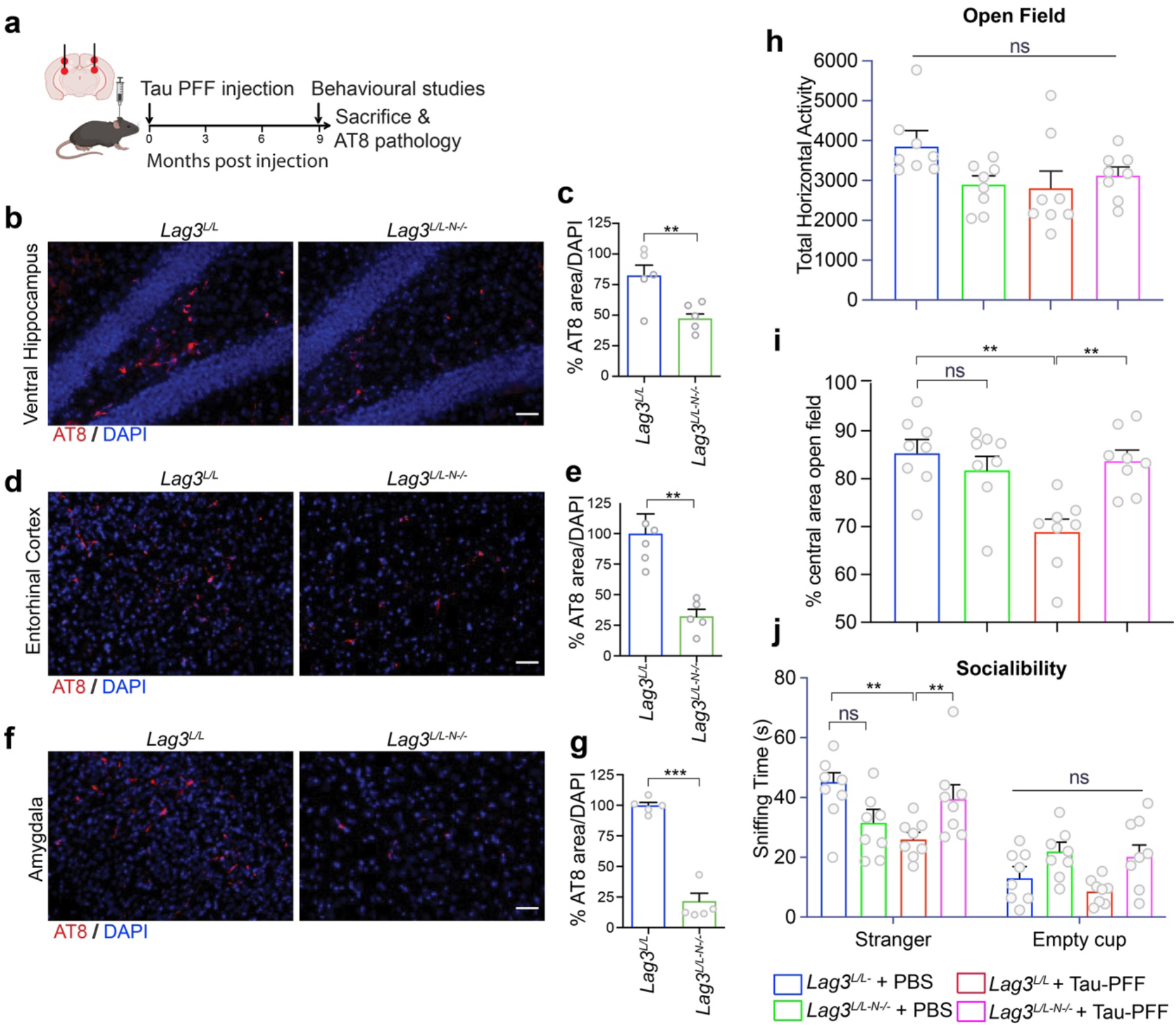
Tau pathologic spreading is reduced in Lag3 neuronal conditional knockout (*Lag3*^L/L-N-/-)^ mice, and reduction of Tau PFF-induced behavioral deficits. **a.** Schema of in vivo experiment. Stereotaxic injection of Tau PFF was performed into the dorsal hippocampus and overlying cortex in *Lag3*^L/L-N-/-^ mice and *Lag3*^L/L^ mice. Pathology and behavioral assessment were performed 9 months after Tau PFF injection. **b-g.** Immunostaining of P-Tau in the ventral Hippocampus (b-c), entorhinal cortex (d-e) and amygdala (f-g) nine months after Tau PFF injection. Error bars represent SEM, *p*-values were calculated with two-tailed paired student *t*-test. \*\**P* < 0.01, \*\*\**P* < 0.001, *n* = 5 mice, Scale bar, 100 μm h-j. *Lag3*^L/L-N-/-^ reduces the behavioral deficits in the open-field test, total horizontal activity (**h**), % central area (**i**) and sniffing time in social interaction test (**j**). Error bars represent SEM, and *P* values were calculated with one-way ANOVA followed by Tukey’s correction . ns, not significant, \*\**P* < 0.01, \*\*\**P* < 0.001.

To determine the role of neuronal Lag3 in mediating behavioral deficits induced by Tau PFF, we performed several behavioral tests as described^33^. We assessed anxiety-related behavior by analyzing time spent in the center zone versus peripheral zone using the open field test^34^. The results show that there is no significant difference in total horizontal activity in the *Lag3*^L/L-YFP^ mice between the PBS and Tau PFF inoculated groups (Fig. 5h), which is consistent with a prior study^33^ indicating that hippocampal injection of Tau PFF cannot cause motor dysfunction. We further assessed the staying time in the central area which reflects anxiety. The results show that Tau PFF can significantly reduce the staying time in the central area in *Lag3*^L/L-YFP^ mice compared to the PBS group (Fig. 5i). In contrast, *Lag3*^L/L-N-/-^ mice alleviated the behavioral deficit in the Tau PFF group (Fig. 5i). Further, we assessed cognitive impairment in the form of sociability with the three chamber social interaction test. The results show that PBS treated mice exhibited normal sociability. In contrast, Tau PFF treated *Lag3*^L/L-YFP^ mice exhibited significantly reduced sociability (Fig. 5j, Extended Data Fig. 8a). As expected depletion of neuronal Lag3 significantly recovered social ability in the Tau PFF group, and there is no significant difference compared to PBS group (Fig. 5j). To evaluate working memory, we tested Tau PFF injected *Lag3*^L/L-YFP^ and *Lag3*^L/L-N-/-^ mice in the Y-maze. We did not find significant behavioral deficits in the Y-maze test by Tau PFF in either group, which is consistent with a prior study^33^ (Extended Data Fig. 8b-d).

## Conclusion

The major finding of this paper is that Lag3 is a Tau PFF-specific receptor, which is required for neuronal uptake and transmission. Although Tau has other receptors, Lag is a receptor that specific for fibrillar Tau. Using Lag3 antibodies and genetic depletion of *Lag3*, Tau PFF-induced pathology and subsequent transmission was reduced. Furthermore, both *Lag3*^-/-^ neurons and Lag3 neuronal conditional knockout mice exhibit significantly reduced spread of Tau pathology and Lag3 neuronal conditional knockout mice had reduced behavioral deficits (Fig. 6).

**Figure 6:**
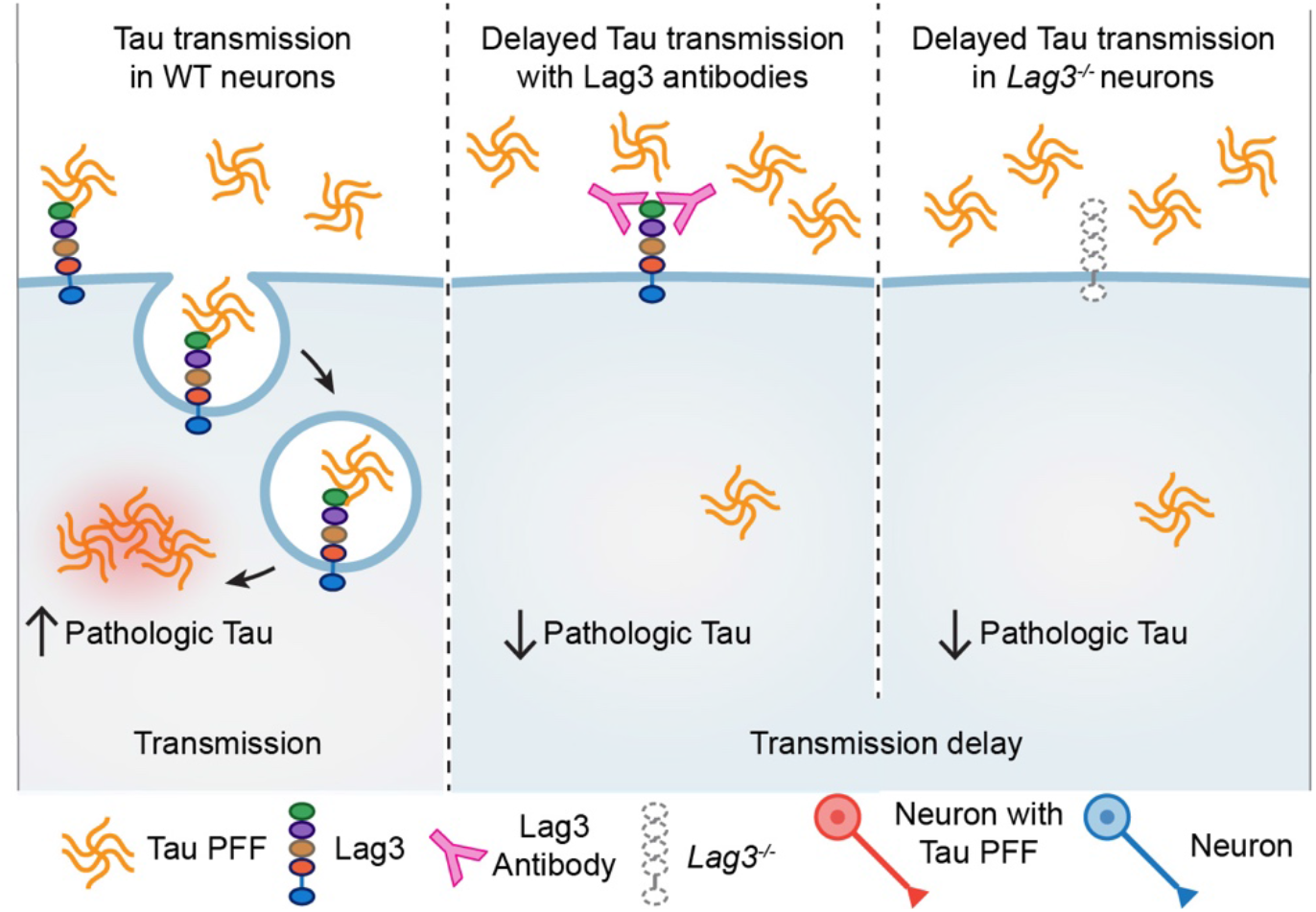
Proposed Model. Conditional Lag3 deletion or antibody-dependent inhibition delays Tau PFF transmission: Tau PFF binding to Lag3 promotes transmission. Binding and endocytosis of Tau PFF are significantly reduced upon *Lag3*^-/-^ or binding to Lag3 antibodies, leading to delayed pathological transmission and toxicity.

LRP1 belongs to low-density lipoprotein receptors (LDLRs) that works in conjunction with HSPGs^35^. HSPGs and LRP1 have been identified as the receptors that bind Tau monomer, oligomer and PFF^9–13^. Deletion of LRP1 can block the uptake and transmission of soluble non-aggregated Tau in the adeno-associated virus (AAV)-transduced WT or P301L Tau models of Tau transmission^8^. The uptake of soluble Tau seems to occur via macropinocytosis^36^, but the neuronal uptake of pathogenic Tau aggregates may rely on endocytosis. In this study, we show that Tau fibrils, but not monomer of Tau can specifically bind to Lag3, and that neuronal Lag3 is required for the endocytosis of Tau PFF. Emerging studies have shown the expression of these receptors are on neurons, microglia, and astroglia where they exert diverse functions^12, 13, 15–18, 37–41^. It would be valuable to determine the roles of these Tau receptors in different cell types.

Heparin-induced Tau fibrils were thought to resemble AD paired helical filaments (PHFs)^42–44^ with strong seeding ability in cellular and animal models overexpressing human mutant Tau; however, hep-Tau PFF can only induce low levels of Tau pathology in non-transgenic mice^3, 7^, which is likely due to different structures^45^. Consistent with this notion in the observation that the structures of hep-Tau PFF are polymorphic and differ from those in AD patients^45^. Our previous studies failed to see the interaction between hep-Tau PFF and Lag3^15, 16^.

These results indicate that the conformation of AD-relevant Tau seeds is critical for prion-like propagation, and the Tau PFF-specific receptors (i.e., Lag3) that recognize the pathogenic conformation are required. It is noted that different Tau species (oligomer, PFF, and large fibrils) and strains can cause different spreading patterns^3, 4, 46, 47^. It will be interesting to further study the interaction between these receptors and Tau species/strains.

The interaction between Lag3 and Tau PFF provides a new target for therapeutic development in AD and related Tauopathies. Repurposing LAG3-blocking antibody combination therapy has been approved by the US FDA with PD-1 inhibition^48^ could be used for neurodegenerative diseases characterized by pathologic tau and α-syn^15, 17, 18^..

## Materials and Methods

### Recombinant Tau monomer purification and endotoxin removal

Recombinant Tau was purified as previously described^3^. WT human Tau isoform (T40, 2N4R) with 441 amino acid residues was cloned into the bacterial expression vector pRK172 provided by Dr. Virginia Lee^3^. Tau was produced in *E. coli* BL21(DE3) strain (Invitrogen, Grand Island, NY, USA). Bacteria were grown in LB broth containing ampicillin at 37 °C. IPTG (Isopropyl β-D-1-thiogalactopyranoside) (1 mM) was used to induce protein generation when the OD reached 0.8 to 1. After 2 hr induction, bacteria were pelleted by centrifugation. High-salt RAB buffer [0.1 M MES, 1 mM ethylene glycol-bis(2-aminoethylether)-N,N,N’,N’-tetraacetic acid (EGTA), 0.5 mM MgSO4, 0.75 M NaCl, 0.02 M NaF, 1 mM phenylmethylsulfonyl fluoride (PMSF), 0.1 % protease inhibitor cocktail (10 ug/mL each of pepstatin A, leupeptin, tosyl phenylalanyl chloromethyl ketone (TPCK), tosyl-L-lysyl-chloromethane hydrochloride (TLCK), and soybean trypsin inhibitor and 100 mM ethylenediaminetetraacetic acid (EDTA)), pH 7.0] was used to resuspend the pellet and followed incubation on ice for 1 hr after homogenization. Then, cell lysates were boiled for 15 min, followed by rapid cooling (on ice for 30 min) and centrifuging (at 30,000g for 30 min). Supernatants were harvested and dialyzed into buffer A [20mM piperazine-N,N′-bis(2-ethanesulfonic acid), 10 mM NaCl, 1 mM EGTA, 1 mM MgSO4, 2 mM dithiothreitol (DTT), 0.1 mM PMSF, pH 6.5], then separated by fast protein liquid chromatography system using HiTrap SP HP column (GE Healthcare, Life Sciences, Wauwatosa, WI, USA). Sodium dodecyl sulfate-polyacrylamide gel electrophoresis (SDS-PAGE) followed by Coomassie Blue R-250 staining was performed to verify the presence and purity of Tau protein. Proteins were concentrated and exchanged with phosphate-buffered saline (PBS) pH7.4 using an Amicon YM-10 centrifuge concentrator (Millipore, Bed-ford, MA).

Pierce high-capacity endotoxin removal spin 0.5 ml column (Thermo Fisher Scientific, Waltham, MA, USA) was used for endotoxin removal. Briefly, 5 ml of ultrafiltrated phage lysate in SM buffer supplemented with 400 mM, NaCl, or 2 ml of AEX-purified and ultrafiltrated phage in the same buffer were applied in the column and incubated at 4 °C for 1 hr. Approximately 4.8 ml of the phage solution was collected by centrifuged at 500g at room temperature (RT) for 1 min. Next, 0.2 N NaOH in 95% ethanol was used to regenerate the column for 1-2 hr at RT. After centrifuging at 500 g for 1 min, the solution was discarded and followed by resuspension of resin with 2 M NaCl and endotoxin-free ultrapure water separately. Then PBS pH 7.4 (endotoxin free) was used to suspend the resin for a total of 3 times. Purified Tau monomer protein (around 10 mg) was loaded into the column and incubated at 4 °C for 1 hr, followed by collection through centrifuge at 500 g for 1 min. Quantitation of endotoxin levels in Tau protein after endotoxin removal was measured by using Pierce™ chromogenic endotoxin quant kit (Thermo Fisher Scientific, Waltham, MA, USA).

### Preparation and characterization of Tau PFF

In vitro, fibrillization of purified recombinant Tau was prepared by mixing 2 mM DTT with 300 µM recombinant Tau in 100 mM sodium acetate buffer (pH 7.0) under constant orbital agitation (1,000 rpm) at 37 °C for 7 days^49^. Successful fibrillization was confirmed using the thioflavin T fluorescence assay and transmission electron microscopy. The fibrils were mechanically broken down into small fragments by sonication (60 sec, ∼0.5 sec/pulse, 10% amplitude). Sulfo-NHS-LC-Biotin was used to label Tau as described previously^15^.

For the extraction of Tau aggregates from AD brains, human brain tissues from two AD patients were were obtained from the Division of Neuropathology, Department of Pathology at Johns Hopkins Schoolof Medicine. Tissue was kept at -80°C. The diagnosis of both cases was histologically confirmed. The purification of AD paired helical filaments (PHF), AD-Tau, was based on a previously published protocol^3^. Briefly, nine volumes (v/w) of high-salt buffer (10 mM Tris-HCl, pH 7.4, 0.8 M NaCl, 1 mM EDTA, and 2 mM DTT, with protease inhibitor cocktail, a phosphatase inhibitor, and PMSF) with 0.1% sarkosyl and 10% sucrose were added to about 2-4 g frozen frontal cortical gray matter, followed by homogenization and centrifugation at 10,000 g for 10 min at 4 °C. The same buffer was used to re-extract the pellets two times. All the three initial extractions were pooled and filtered, and additional sarkosyl was added to reach 1 %. The collected supernatant was centrifuged (300,000 g for 60 min at 4°C) again after 1 h nutation at room temperature. The pellets (1% sarkosyl-insoluble) containing pathological Tau were washed once in PBS, resuspended, and passed through a 0.5mL TB Syringe with 27 Gauge 1/2 inch permanently attached needles in PBS (∼100 µl/g gray matter). Further purification through brief sonication (20 pulses at ∼0.5 sec/pulse) and centrifugation (100,000 g for 30 min at 4 °C) was used for the resuspended sarkosyl-insoluble pellets. Then, the pellets were resuspended in PBS at one half of the precentrifugation volume, sonicated with 20–60 pulses (∼0.5 sec/pulse), boiled for 10 min and spun at 10,000 g for 30 min at 4 °C to remove large debris. The final supernatants were collected as the final enriched AD extracts.

The in vitro fibrillization of recombinant Tau by AD-Tau seeds was performed following a previous protocol^3^. Briefly, 4 µM AD-Tau was used as seeds for a total concentration of 36 µM Tau in DPBS (endotoxin free), and the mixture was kept on an agitator for 3-4 days (1000 rpm, 37 °C). The pellets were collected after centrifugation (100,000 g for 30 min at 4 °C), and washed once in DPBS (endotoxin free) and resuspended in DPBS (endotoxin free), which were aliquoted and stored at -80 °C referred to as Tau PFF. Right before each use, the aliquots were briefly sonicated with 30 pulses (∼0.5 sec/pulse) and spun at 10,000 g for 15 min at 4 °C to remove large debris.

### Transmission electron microscopy (TEM) measurements

Negative stain electron microscopy was used to verify the formation of fibrils for Tau PFF. Briefly, formvar carbon coated TEM grids were drop casted with 10 µL of Tau PFF for 2 min at room temperature, rinsed with water and excess water soaked with lint-free filter paper, and then stained with 2% (w/v) uranyl acetate for another 1 min. The stain was blotted off with lint free filter paper. Grids were allowed to dry completely before taking images captured on Hitachi TEM operating at 80 kV.

### Primary neuronal cultures, Tau PFF administration and neuron binding assays

Primary cortical neurons were prepared from E15.5 pups of C57BL/6 mice and cultured in neurobasal media adding with B-27, 0.5 mM L-glutamine, streptomycin, and penicillin (Corning, glendale, AZ, USA) on tissue culture plate coated with poly-L-lysine. The medium was changed every 3 days. At 7 days in vitro (DIV), Tau PFF (2 µg/mL) was added and incubated for 14 days in the medium. Subsequently, biochemical experiments and pathology assays were performed. Tau-biotin PFF at different concentrations were utilized to define the amount of bound Tau-biotin PFF to WT and *Lag3*^−/−^ neurons. ImageJ software was used for the quantitative analysis.

### SH-SY5Y cell surface binding assays

SH-SY5Y cells were transfected with Lag3 expressing plasmid with Lipofectamine 2000 as per the manufacturer’s instruction. Transfected cells were treated with different concentrations of monomer equivalent Tau-biotin PFF in DMEM media (10% FBS) for 2 hr at RT. We removed unbound Tau-biotin PFF by thoroughly washing (5 times, 20 min each) with DMEM media (10% FBS) on a horizontal shaker, followed by fixation with 4% paraformaldehyde in PBS. The cells were washed three times with PBS, followed by blocking with PBS containing 10 % horse serum and 0.1% Triton X-100 for 30 min. After blocking, the cells were incubated with alkaline-phosphatase-conjugated streptavidin (1:2000 dilution) in PBS with 5 % horse serum and 0.05 % Triton X-100 for 16 hr. Alkaline phosphatase signal was detected histochemically by 5-bromo-4-chloro-3-indolyl phosphate (BCIP)/nitro blue tetrazolium (NBT) reaction^19–21^. Images were acquired on Zeiss Axiovert 200M microscope. The intensity of bound Tau-biotin PFF to *Lag3*-transfected SH-SY5Y cells was quantified with ImageJ software. Desired range of intensity values and background exclusion for control was acquired by adjusting the threshold under the Image/Adjust menu. The same threshold settings were applied to all images in each experiment. The binding curve and KD values were calculated with GraphPad Prism software.

### Plasmids and deletion mutants

Human LAG3 and mouse *Lag3* cDNA clones were kindly obtained from Dr. Charles Drake at the Johns Hopkins University School of Medicine. Lag3 deletion mutants (D1, D2, D3, D4, TM, and ICD) with a HA tag and deletion mutants of Lag3 D1 domain with a myc tag were constructed by PCR with In-Fusion HD Cloning Plus systems (Takara; Bio Inc., Otsu, Japan) using the CloneAmp™ HiFi PCR Premix (a high-fidelity PCR polymerase included with all In-Fusion HD Cloning Plus Systems), and primers with a 15 bp overlap with each other at their 5’ ends and incorporate the mutation of interest. The DNA was separated on a 1% agarose gel and the appropriate band was excised and isolated using a gel extraction kit (Qiagen, Valencia, CA, USA). In-Fusion HD enzyme premix was added to the linearized PCR product and transformed into the Stellar™ Competent Cells (Takara; Bio Inc., Otsu, Japan). Integrity of plasmid sequences was verified through sequencing.

### Lentiviral vector construction, production and transduction

Preparation of lentiviral vectors was performed based on a prior publication^50^. Briefly, *Lag3* or *Lag3* deleted mutations with HA tag were subcloned into a lentiviral cFugw vector through Age I restriction enzyme site. The human ubiquitin C (hUBC) promoter was utilized to drive gene expression. The recombinant cFugw vector gathering with three packaging vectors (pLP1, pLP2, and pVSV-G (1.3:1.5:1:1.5)) was transiently transfected into HEK293FT cells and the lentiviruses were produced. The viral supernatants were concentrated and collected through ultracentrifugation (50,000g, 3 hr) after 48 and 72 hr transfection. Viral particles were stored in serum-free medium at −80 °C. Neurons (DIV 4 to 5) were transducted by lentivirus separately carrying Lag3, Lag3 deletion mutants for 72 hours, while empty vectors served as a control [1 × 109 transduction units (TU)/ml].

### Live images

Primary cultured neurons were cultured on 12-mm coverslip in a chamber (RC-48LP, Warner Instruments) with physiological saline solution (40 mM NaCl, 5.4 mM KCl, 1.3 CaCl_2_ mM, 1.0 mM MgCl_2_, 20 mM glucose, and 25 mM Hydroxyethyl piperazine Ethane Sulfonicacid (HEPES), pH to 7.4) and imaged using Microscope Axio Observer Z1 (Zeiss, Dublin, CA, USA). Tau PFF was labeled with pHrodo red (Invitrogen, Grand Island, NY, USA). pHrodo red fluorescence increases as the pH drops when it gets into neurons. Tau PFF-pHrodo was directly added to Lag3 WT and *Lag3*^−/−^ neuron groups. For the WT + Lag3 and *Lag3*^−/−^ + Lag3 groups, neurons were transduced with lentivirus carrying *Lag3* (empty vector as a control). Live images were recorded every 0.5∼1 min for 90∼100 min. 1∼4 min later after 0.5 µM Tau PFF-pHrodo treatment, the cells were appropriate for recording. Fluorescence intensity of the neuron at 2–3 min after Tau PFF-pHrodo treatment was used as the baseline background. Quantification was performed using the Zeiss Zen Software to outline the neurons and subtract the background. The percentage of internalized Tau PFF-pHrodo signal at each time point was calculated based on the ratio to the baseline in each experiment.

### Calcium imaging

Calcium imaging was utilized to visualize calcium signaling in a living neuron directly through intracellular calcium flux. In our study, we used Fluo-2 acetoxymethyl (AM) ester (Thermo Fisher Scientific, Waltham, MA, USA) as the fluorescent calcium indicator to monitor the intracellular calcium levels in primary cultured cortical neurons. Primary cortical neurons obtained from WT and *Lag3*^−/−^ embryo mouse cortex were plated on coverslips coated with polyornithine and incubated for 14 days. Right before imaging, the cells were treated with Fluo-2 AM for 30 min at 1 µM final concentration. Fura-2-free PSS buffer was used to wash the cells; then the neurons were put in a chamber (RC-48LP, Warner Instruments) with a 37°C heated adaptor on the Microscope Axio Observer Z1 (Zeiss, Dublin, CA, USA). The field containing more than 10 neurons was selected as Regions of interest (ROI). The imaging was conducted with an excitation/emission wavelength of 485 nm/525 nm. After the baseline fluorescent signals were kept steady for 5 min, 500 nM Tau PFF was added and the imaging recording lasted for another 90 min. During the imaging recording process, images were captured every 30 sec and analyzed by Image J and Zen software (Carl Zeiss, Germany).

### Mouse strains

C57BL/6 wild-type (WT) (strain 000664) mice were purchased from the Jackson Laboratories (Bar Harbor, ME), while *Lag3*^−/−^ mice were kindly provided by Dr. Charles Drake (Johns Hopkins University School of Medicine) maintained on a C57BL6 background. *Lag3*^L/L-YFP^ mice have been previously described^30^. Nestin^Cre^ mice^31, 32^ were obtained from Jax lab (strain 003771). Thesed mice did not develop any autoimmune or inflammatory phenotype. The research was approved by the Johns Hopkins University Animal Care and Use Committee and all procedures were completed according to the NIH Guide for the Care and Use of Experimental Animals (NIH Publication No. 85-23, revised 1996). Animals were housed with free access to water and food at a 12:12 hr dark–light cycle. Before performing any animal experiments, all mice were kept in the procedure room for an acclimatization period.

### Lag3 antibody blocking experiments

Anti-Lag3 antibodies (410C9, C9B7W, 17B4) were added to the cell cultures (30 nM) 30 min before the Tau PFF treatment, while the rat IgG (Invitrogen, Carlsbad, CA, USA) and mouse IgG (Santa Cruz, Dallas, TX, USA) were used as negative controls. We used SH-SY5Y cells that overexpressed Lag3 for the binding assay two days after Lag3 transfection. For the endocytosis endosome enrichment assay, mouse primary cortical neurons were cultured for 12 days (12 DIV) before treatment. For Tau pathology and transmission experiments, Tau PFF was added to wells of primary cultured neurons cultured for seven days in vitro (7 DIV). In the neuron transmission experiment, the antibodies were added to the middle chamber (chamber 2) of the microfluidic chamber system.

### Microfluidic chambers

Triple compartment microfluidic devices (TCND1000, Xona Microfluidic, LLC, Temecula, CA, USA) were used for Tau transmission experiment in vitro. After the glass coverslips were prepared and coated as previously described, they were attached to the microfluidic device^51^. About 100,000 primary cultured neurons were plated in each chamber of the device. Lentivirus *Lag3* was used to transduce WT and *Lag3*^−/−^ neurons (4 DIV) to create WT + Lag3 and *Lag3*^−/−^ + Lag3 neurons. Then, Tau PFF (1 µg/ml) was added into chamber 1 at 7 DIV. In order to control the direction of medium flow, the difference of medium volume (between chamber 1 and chamber 2, and between chamber 2 and chamber 3), was maintained at 75 µL based on the manufacturer’s instructions. After 14 days of treatment, neurons were fixed using 4 % paraformaldehyde in PBS, followed by immunofluorescence staining.

### Stereotaxic injection procedure

Aliquoted Tau PFF was used in this experiment. Right before stereotaxic injection, the Tau PFF was diluted with PBS and briefly sonicated in a temperate controlled water bath sonicator (∼0.5 sec/pulse, 30 pulses) and spun at 10,000 g for 15 min at 4 °C to remove large debris. Pentobarbital was used for the anesthesia of 3 months old C57BL6 mice. Tau PFF (2 µg/2 µL) was delivered into two sides of the dorsal hippocampus and overlaying cortex using the stereotactic apparatus at reference coordinates for the hippocampus (Bregma −2.54 mm, 2 mm from midline, and −2.4 mm from skull) and the cortex (Bregma −2.54 mm, 2 mm from midline, and −1.4 mm from skull) based on previous publication^3^. Hamilton syringe (2 µL, Hamilton, Reno, NV, USA) was used for the injections with a rate of 0.1 µL/min, and the needle was kept in place for ≥ 5 min before gentle removal. Animals were carefully monitored and taken care of after the operation. Animal behavior tests were performed at 9 months after injection. After euthanization, the mice were used for neurochemical, biochemical and histological studies. Tissues were immediately frozen after removal and stored at −80 °C for biochemical experiments. For immunofluorescent staining studies, mice were perfused transcardially with PBS followed by fixation with 4% Paraformaldehyde (PFA), and brains were removed and fixed overnight in 4 % PFA, followed by keeping in 30% sucrose in PBS until tissue settled down.

### Immunofluorescence and mapping of Tau pathology

Serial brain sections (40 µm thick) were used to perform immunohistochemistry (IHC) and immunohistofluorescence (IHF). Coronal sections were incubated in primary antibodies for P-Tau (AT8) (Invitrogen, Grand Island, NY, USA) Immunoreactivity was labeled using appropriate fluorescent secondary antibodies conjugated to Alexa-fluor 568 (Invitrogen, Carlsbad, CA, USA). DAPI was used together with secondary antibodies to stain the nuclei. Images (IF) were detected by confocal scanning microscopy (LSM 880, Zeiss, Dublin, CA, USA) or fluorescent microscope (BZ-X710, Keyence, Osaka, Japan).

### Behavioral analysis

Behavior tests, such as open field, Y-maze and social interaction tests were used to assess behavioral deficits in mice at 9 months after the injection of PBS control or Tau PFF. The experiment was performed with a blinded treatment group for all behavioral tests. All tests were performed between 13:00–17:00 in the lights-on cycle.

### Open field test

General motor activity and anxiety-related behavior were assessed by an open field test. In brief, we randomized all mice by four open field chambers to mix up control versus experimental groups. Then, the mice were directly placed into the middle of the open field (a rectangular plastic box, 40 cm × 40 cm × 40 cm, length × width × height). The plastic box was divided into 36 identical sectors and the field was subdivided into peripheral and central sectors^52^. Movements were recorded for 5 min using a digital camera under dim light. The open field was wiped down with Vimoba after the removal of each experimental mouse. Distance traveled was recorded using the Photobeam Activity System for the analysis of locomotor activity, and the percentage of time spent in the center square was measured for anxiety analysis.

### Y-maze

We used Y-maze to evaluate the spatial working memory of mice. The experiment consisted of the training and test phases were performed as previously described^53^. Briefly, the mice were collected from their rooms and brought to the behavioral suite to allow for acclimation. The Y-maze apparatus consists of three symmetrical arms with an angle of 120° between adjacent arms. (15 cm x 10 cm x 40 cm, height × width × length, each arm A, B, C). The three arms were randomly allocated for the start arm, familiar arm, and novel arm. During the training phase, mice were allowed to explore start arm and familiar arm freely for 8 min while the novel arm was blocked by a removable barrier. After the training phase, the animal was removed from the Y-maze apparatus and kept in their home cage for 2 hr. Then the test was conducted while the barrier wall of the novel arm was removed. The mouse had access to all three arms freely for 8 min, starting at the same end of the start arm. Behavioral measurements were performed and analyzed by Any-Maze software. Additionally, we also recorded a spontaneous alternation when the mouse entered the three arms in turn, such as CBA, ABC or BAC. The time spent in each arm, the number of arms entered and the number of alternations or triads were recorded. In addition, the relative time spent in the novel arm and the number of alternations were calculated. Finally, the total distance the mice moved on the entire maze was recorded and analyzed in order to assess general locomotor activity.

### Social interaction tests

Three chamber social interaction tests were conducted to evaluate the mice’s sociability and social novelty. In the first phase (habituation), we placed the mice into the 3-chambered apparatus without any stranger mice, opened all the gates, and allowed the mouse to explore for 10 min. In the second phase (sociability), we closed the gates of the chambers and kept the test mouse in the center of the apparatus. Subsequently, the “stranger 1” mouse was placed under one of the wire baskets while the other basket with no animal inside was placed in the other chamber. Then, we opened the gates and allowed it to explore for 10 min. In the third phase (preference for social novelty), we closed both gates, placed the “stranger 2” mouse under the empty wire basket in the second phase, and allowed the test mouse to explore for 10 min. The sniffing time at Stranger 1 and Stranger 2 was recorded using Topscan 3.0 software.

### Quantification and Statistical Analysis

All data were analyzed using GraphPad Prism 8. Statistics Data are presented as the mean ± SEM with at least 3 independent experiments. Representative morphological images were obtained from at least 3 experiments with similar results. Statistical significance was assessed via a one or two-way ANOVA test followed by indicated post-hoc multiple comparison analysis. Assessments with *p* < 0.05 were considered significant.

## Acknowledgements

T.M.D. is the Leonard and Madlyn Abramson Professor in Neurodegenerative Diseases. The Multiphoton Imaging Core of Johns Hopkins University was used (NS050274) in some of the imaging studies.

## Author contributions

CC and RK led the project and contributed to all aspects of the study. HW, XLY, KG, CR, YK, AB, SK, DJ, LW, AW, RC, SZ contributed to biochemical, cellular, mouse experiments work. CJW and DAAV provided the essential mice strains and reagents. OP, DWN, JCT provided key brain samples. JLT, PCW and MYY helped with the project/experimental design. XM, VLD, and TMD designed research, XM, VLD, and TMD, wrote and revised the paper. All authors reviewed, edited and approved the paper.

## Data and materials availability

Further information and requests for resources and reagents should be directed to and will be fulfilled by Xiaobo Mao. There are no restrictions on any data or materials presented in this paper. All data are available in the main text or the Extended Data.

## Extended figures and Figure legends

**Extended Figure 1:**
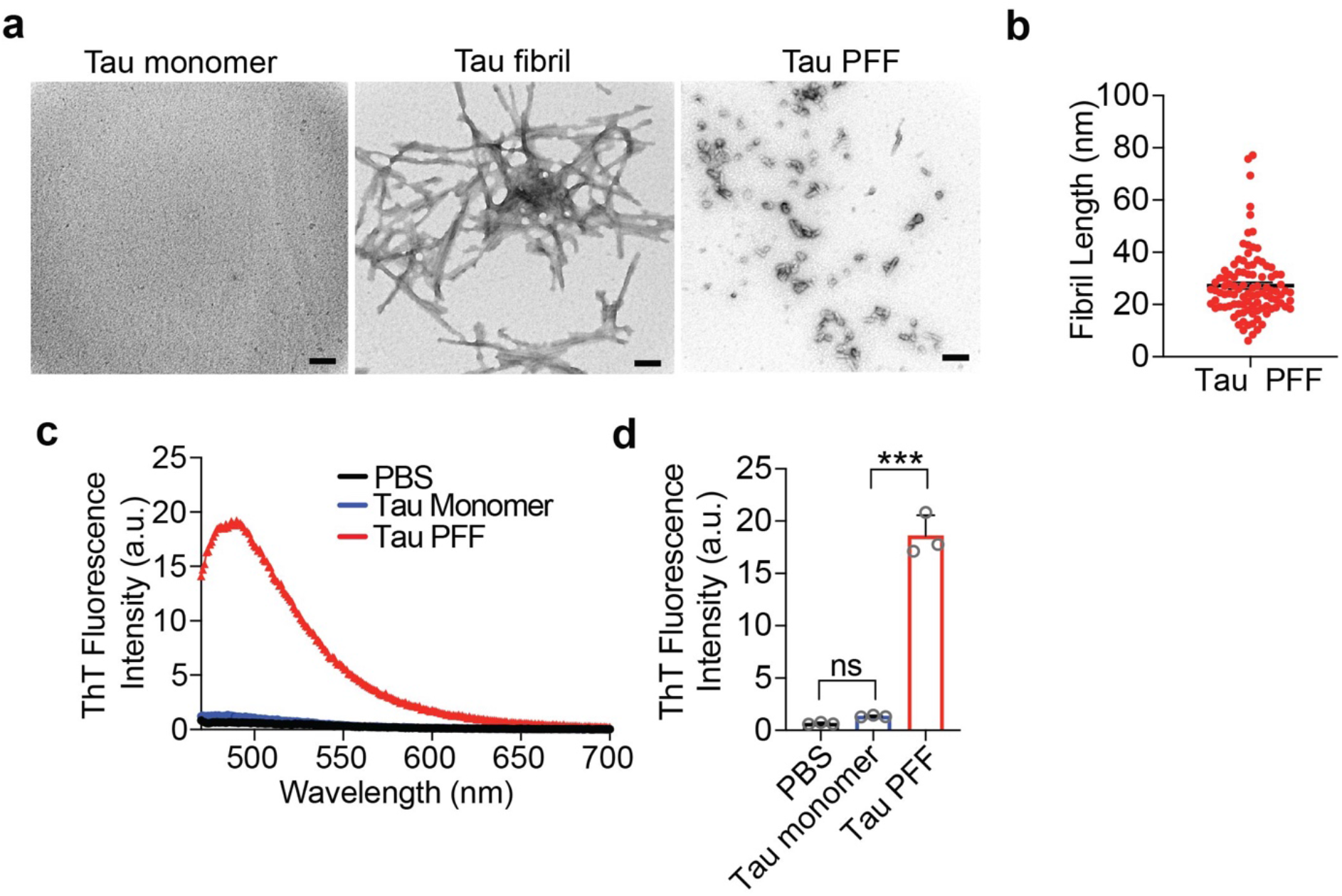
Characterization of Tau PFF. **a**. Transmission electron microscopy of Tau monomer, Tau fibrils (unsonicated) and Tau PFF (sonicated). Scale bar, 100 nm. **b.** Fibril length of sonicated Tau PFF. The error bar represents SEM. **c.** The ThT fluorescence spectra were recorded at 450 nm excitation and 470 to 700 nm emission wavelengths. **d.** ThT fluorescence intensity at 485 nm for PBS, Tau monomer, and Tau PFF (sonicated), (*n* = 3). Error bars represent SEM.

**Extended Figure 2:**
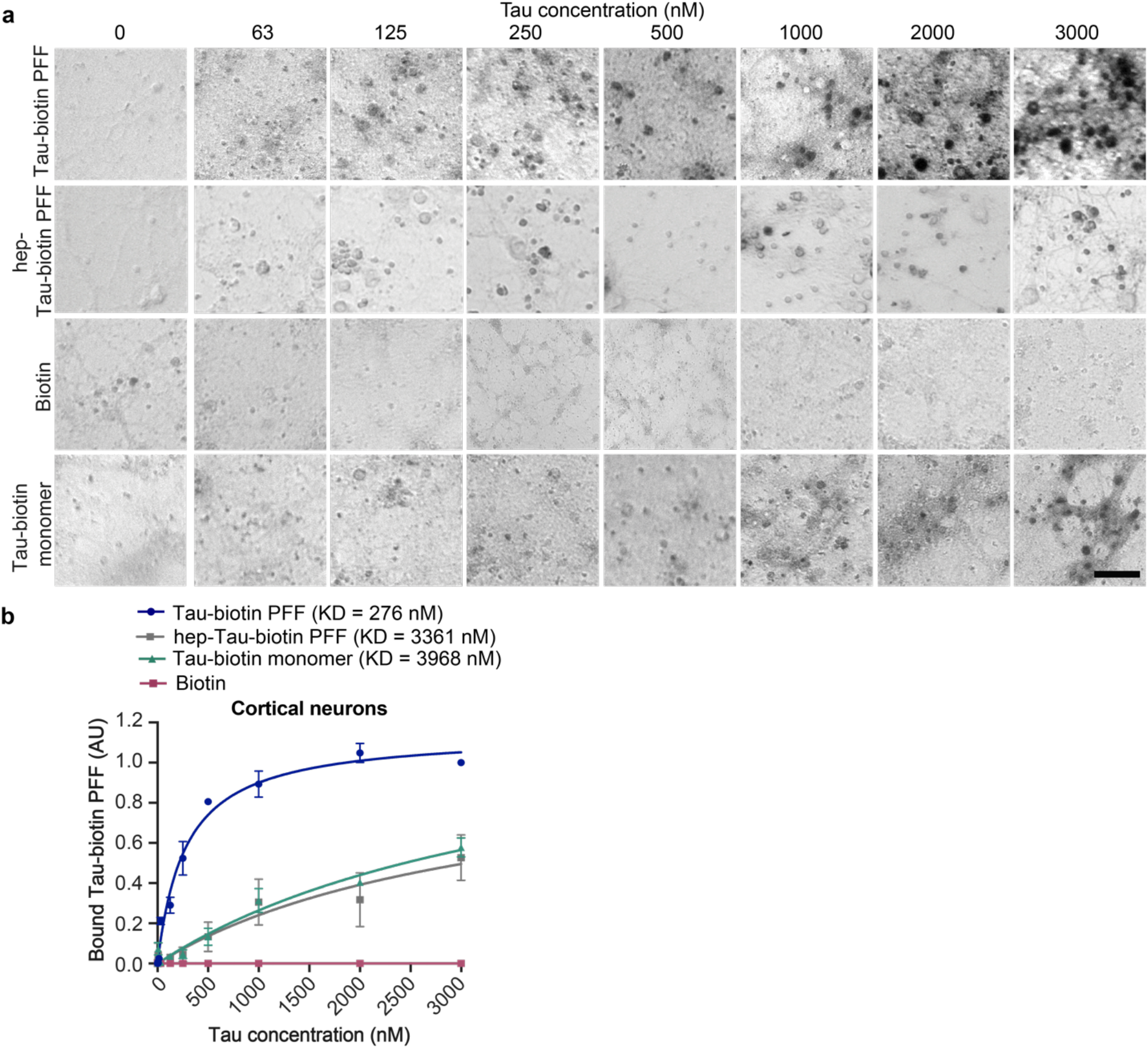
Binding of biotin-labeled Tau PFF to wild-type (WT) and *Lag3*^-/-^ neurons. **a.** Representative images of Tau PFF-biotin binding to wild-type (WT) mouse primary cortical neurons, Scale bar, 100μm. **b.** Quantification of binding of biotin, Tau-biotin monomer and heparin induced Tau-biotin (hep-Tau) PFF and Tau-biotin PFF to mouse primary cortical neurons (Tau-Biotin PFF KD = 276 nM), Data are the means ± SEM, *n* = 3 independent experiments.

**Extended Figure 3:**
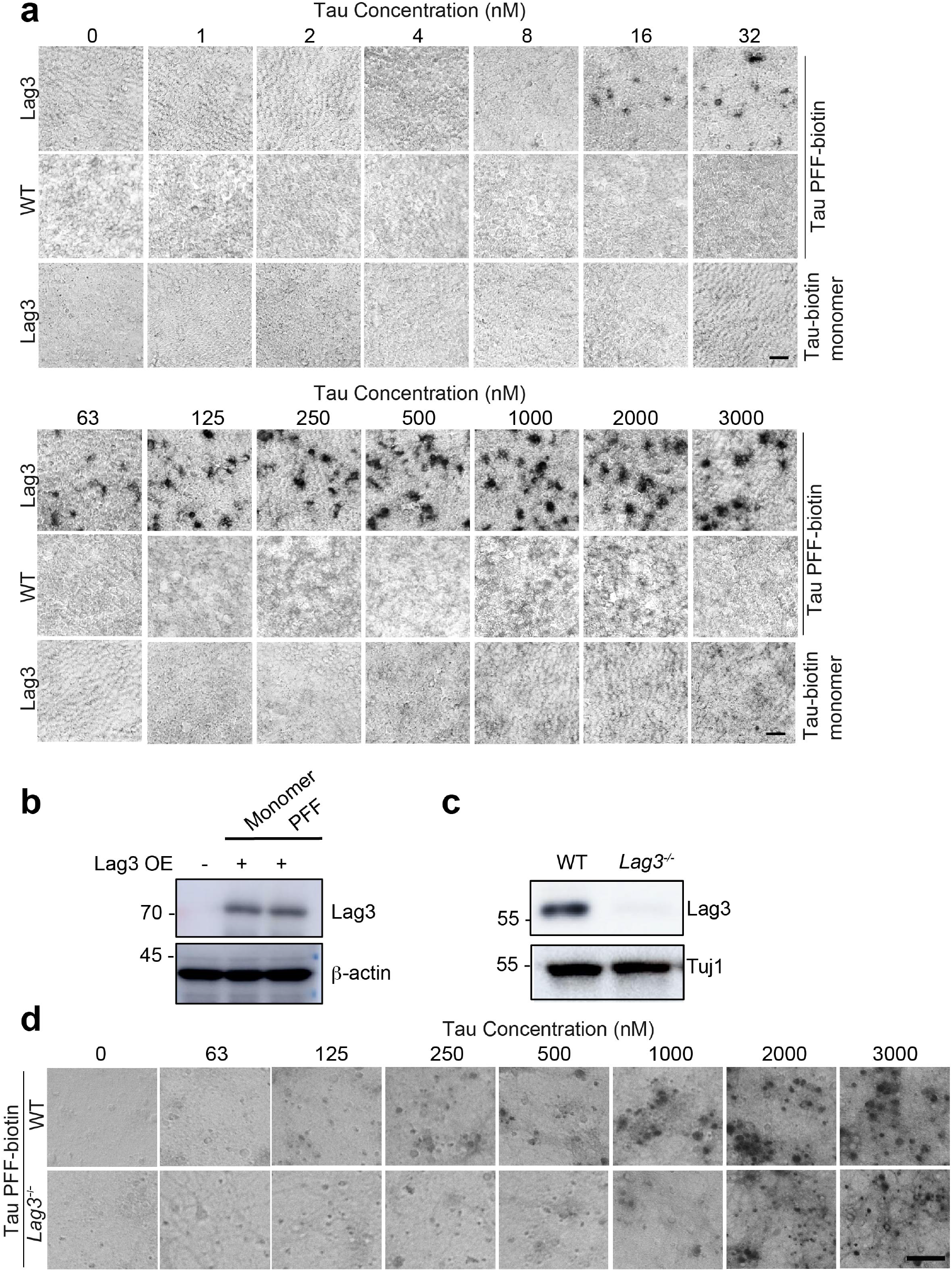
Biotin-labeled Tau PFF can bind to Lag3 expressing SH-SY5Y cell surface receptor. **a.** Tau-biotin PFF binds to Lag3 overexpressing SH-SY5Y cell surface in a saturable manner, as a function of Tau biotin total concentration (monomer equivalent for PFF preparations). Scale bar, 100 μm. **b.** Immunoblot showing overexpression (OE) of Lag3 into SH-SY5Y cells in the cell surface binding assays. **c.** Immunoblot showing Lag3 expression level in WT and *Lag3* knockout (*Lag3*^-/-^) mouse primary cortical neurons. **d.** Representative images of Tau PFF-biotin binding to WT and *Lag3*^-/-^ mouse primary cortical neurons. Scale bar, 100 μm.

**Extended Figure 4:**
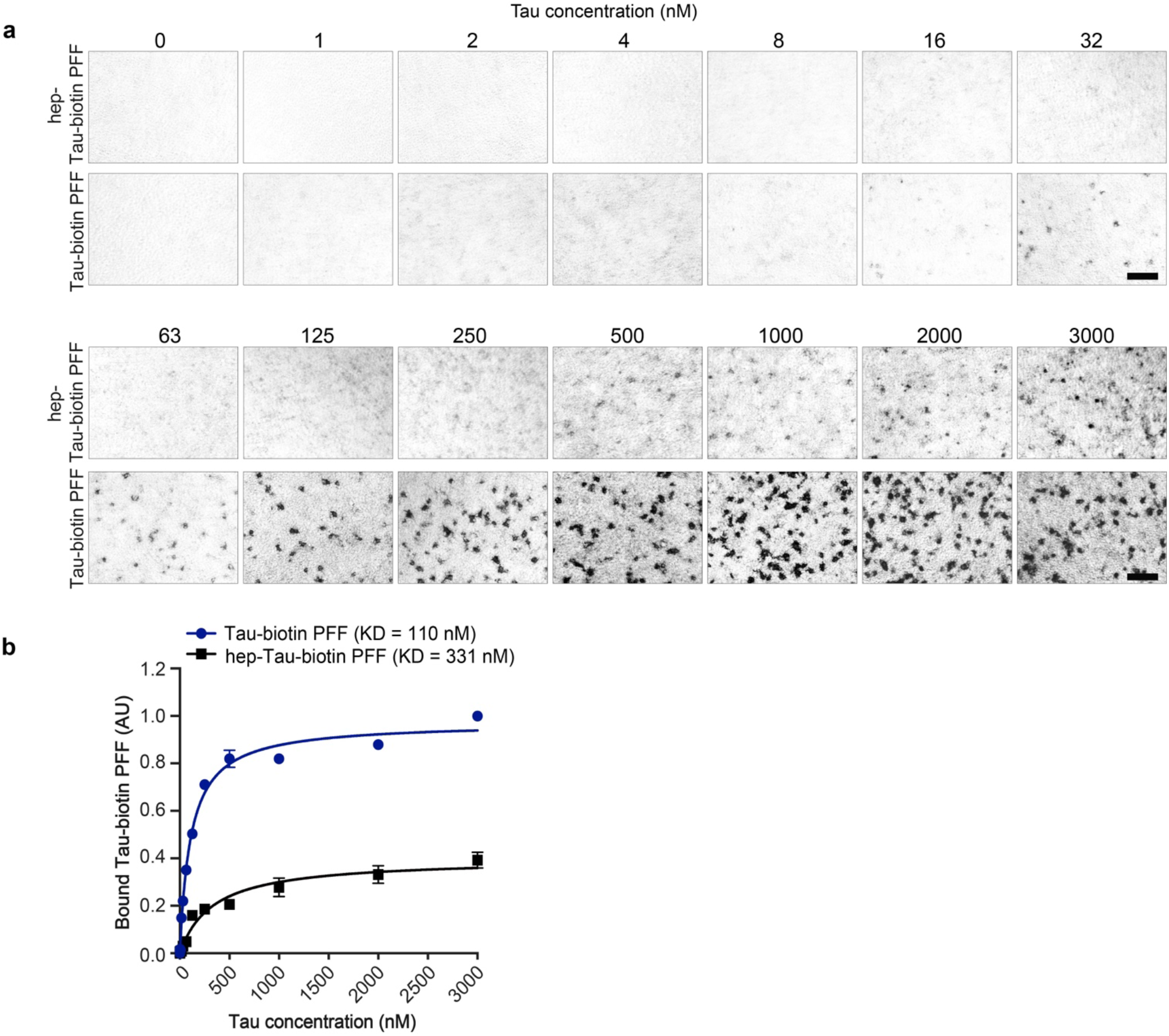
Heparin inhibits Tau PFF binding to Lag3. **a.** Representative images of Tau-biotin PFF and Heparin induced Tau (hep-Tau)-biotin PFF binding to Lag3-expressing SH-SY5Y cell surface. Scale bar, 100 μm. **b.** Quantification of binding of Tau-biotin PFF and hep-Tau PFF-biotin to SH-SY5Y cell surface. Data are the means ± SEM, *n* = 3 independent experiments.

**Extended Figure 5:**
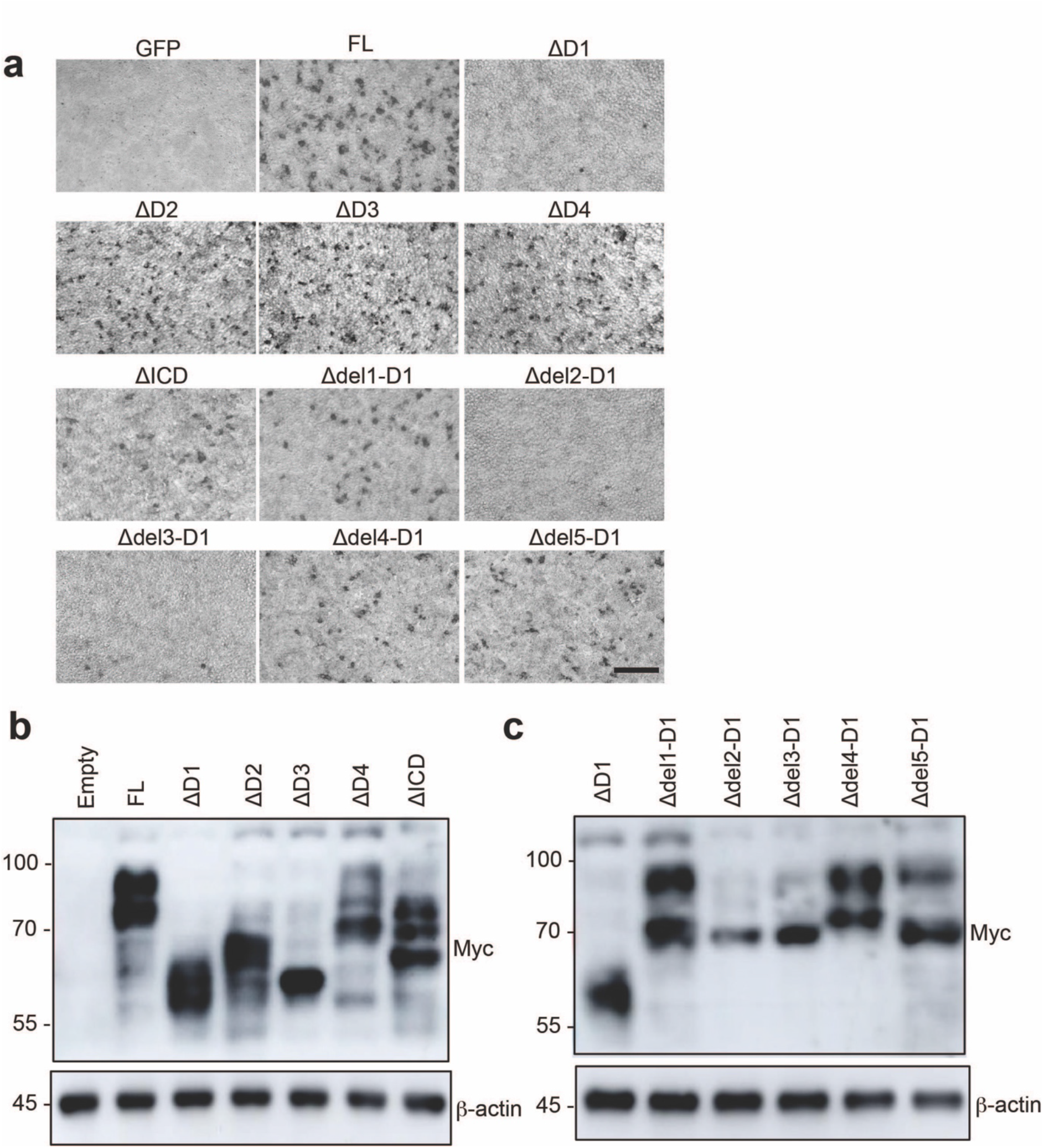
Lag3 deletion mutants were overexpressed into HEK293FT cells and assessed for Tau PFF-biotin binding. **a.** Representative images of Tau PFF-biotin binding to full-length (FL) and deletion mutants of Lag3: extracellular domains (ΔD1-ΔD4), intracellular domain (ΔICD, and subdomains of D1 domain (Δdel1-5-D1). Scale bar, 100 μm. **b-c.** Immunoblot showing overexpression levels of full-length Lag3 and deletion mutants of Lag3: extracellular domains (ΔD1-ΔD4), intracellular domain (ΔICD, and subdomains of D1 domain in HEK293FT cells used for Tau PFF-biotin binding.

**Extended Figure 6:**
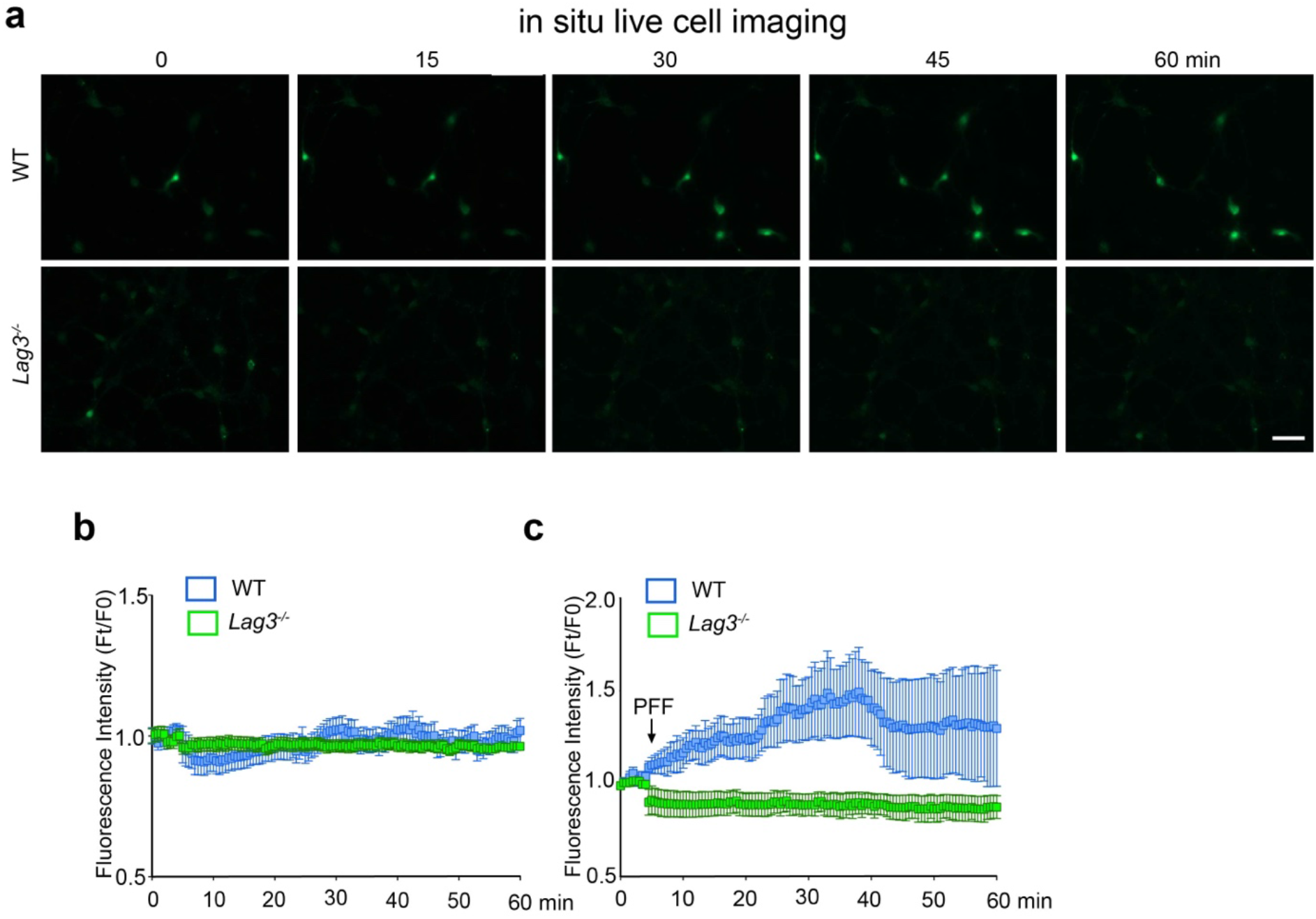
Tau PFF induced increase in intracellular Ca^2+^ level is mediated by Lag3. **a.** Time series imaging of calcium-dependent fluorescence of WT and *Lag3*^-/-^ neurons. Neurons were treated with 1 µM Fluo-2 acetoxymethyl (AM) ester for 30 min followed by 500 nM Tau PFF. Live images were acquired for the indicated time period. Scale bar, 50 µM. **b.** Ca^2+^ levels from WT and *Lag3*^-/-^ neurons without Tau PFF treatment. **c.** Fluo-2 acetoxymethyl (AM) associated fluorometric measurement showing changes in intracellular Ca^2+^ signal in WT and *Lag3*^-/-^ neurons upon Tau PFF treatment. Ft fluorescence intensity of the indicator at t minutes, F0 is Fluorescence intensity at 0 min. Error bars represent SEM.

**Extended Figure 7:**
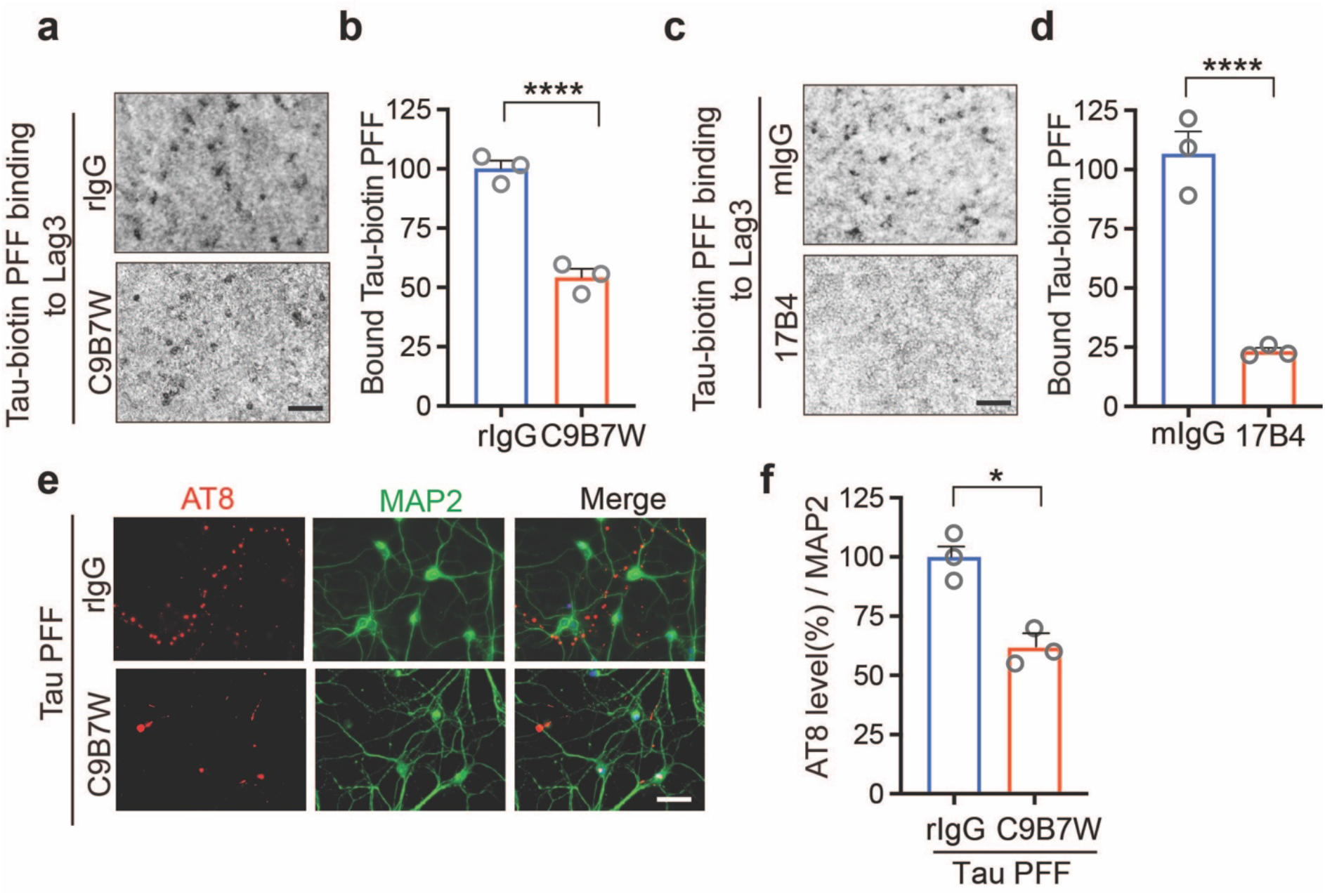
Lag3/LAG3 antibodies block Tau PFF binding to Lag3/LAG3, and subsequent pathologic propagation. **a.** Anti-Lag3 C9B7W blocks the binding of Tau-biotin PFF to Lag3-expressing SH-SY5Y cells. **b.** Quantification of a. Error bars represent means ± SEM, *n* = 3 independent experiments, Student’s *t*-test. \*\*\*\**P* < 0.0001. **c**. Anti-human-LAG3 antibody 17B4 blocks the binding of Tau-biotin PFF to human LAG3-expressing SH-SY5Y cells. **d.** Quantification of c, Error bars represents means ± SEM, *n* = 3 independent experiments, Student’s *t*-test. \*\*\*\**P* < 0.0001.**e.** AT8 phosphorylated Tau (P-Tau) was reduced by treatment with C9B7W in primary cortical neurons. **f.** Quantification of e. Error bars represent means ± SEM, *n* = 3 independent experiments, Student’s *t*-test. \**P* < 0.05, Scale bar, 50 μm.

**Extended Figure 8:**
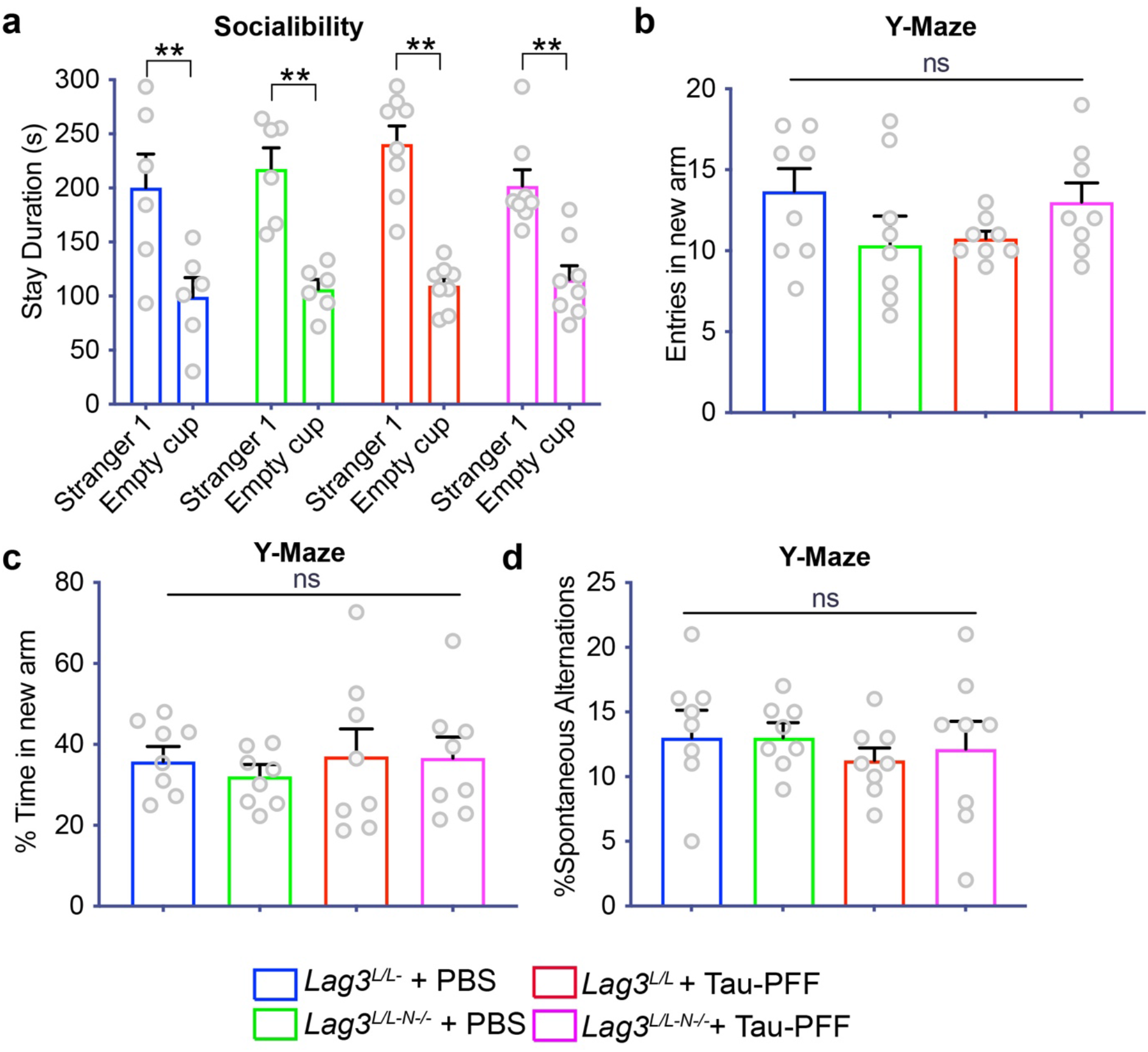
Behavioral assessment of Tau PFF injected *Lag3*^L/L-N-/-^ mice. **a.** Total stay duration in the empty cup and stranger cup in three chamber social interaction test. Student’s *t*-test. \**P* < 0.01, **b-d.** *Lag3*^L/L-N-/-^ mice did not exhibit significant behavioral deficit in the Y-maze test. Specifically, number of entries in the new arm (**b**), percent time spent in the new arm (**c**), percentage of spontaneous alteration of behavior **(d),** Error bars represent SEM, *n* = 8 mice.

